# The inhibition of the JNK2-Syntaxin-1A interaction neuroprotects against retinal degeneration

**DOI:** 10.1101/2025.09.30.679517

**Authors:** Marco Cimino, Jack Serkiz, Joanne K. Konstantopoulos, Annamaria Tisi, Pamela Cappelletti, Rita Maccarone, Rebecca M. Sappington, Marco Feligioni

## Abstract

Retinal diseases (RDs) involve the degeneration of retinal cells, particularly retinal ganglion cells (RGCs), often driven by glutamate imbalance and aberrant signaling. We previously identified a presynaptic self-amplifying mechanism of glutamate overflow, where NMDA overstimulation activates JNK2-mediated phosphorylation of STX1A. To block this mechanism, a cell-permeable peptide, called JGRi1, was previously developed to disrupt the JNK2–STX1A interaction. Here, we investigated whether inhibition of this pathway by JGRi1 could provide neuroprotection in retinal degeneration. Here we showed that JGRi1 efficiently reached the mouse retina upon topical administration as eye drops and granted retinal protection. Using an ex vivo optic nerve cut (evONC) model, we demonstrated that JGRi1 preserved RGC viability, reduced phosphorylation of JNK and STX1A, and lowered glutamate release. In retinal wholemounts, JGRi1 similarly preserved RGC survival. Furthermore, in an NMDA-induced degeneration model, JGRi1 protected RGCs, reduced glutamate levels, disrupted the JNK2–STX1A interaction, and limited microglial infiltration. Collectively, our findings highlight the central role of the JNK2–STX1A pathway in retinal degeneration and identify JGRi1 as a promising neuroprotective tool.

**SYNOPSIS:** Glutamate excitotoxicity drives retinal diseases via JNK2-dependent phosphorylation of STX1A, causing non-canonical presynaptic glutamate spillover (nPING). We developed JGRi1, a cell-permeable peptide that blocks the JNK2–STX1A interaction and prevents spillover.

- JGRi1 effectively reaches the retina via both ex vivo and in vivo topical administration, accumulating in the ganglion cell layer (GCL) and other retinal layers in a dose-dependent manner.
- JGRi1 protects retinal ganglion cells (RGCs) from degeneration in both evONC- and NMDA-induced models, reducing apoptosis, preserving retinal cytoarchitecture and axonal connectivity, and lowering glutamate spillover.
- JGRi1 counteracts the excitotoxic cascade by reducing JNK2 upregulation, STX1A phosphorylation, their interaction and co-localization, limiting SNARE complex formation, and dampening microglial activation.

## INTRODUCTION

Glutamate is the principal excitatory neurotransmitter in the central nervous system (CNS), where it mediates the majority of synaptic transmission. Under physiological conditions, glutamate signaling regulates neuronal communication and plasticity, including mechanisms of learning and memory, primarily via ionotropic and metabotropic glutamate receptors(Zhou e Danbolt, 2014; Andersen et al., 2021). Conversely, dysregulation of glutamate homeostasis leads to excitotoxicity, a condition characterized by excessive activation of glutamate receptors, including N-methyl-D-aspartate (NMDA) receptors (NMDARs), calcium overload, and neuronal injury (Choi, 1992). Excitotoxicity has been implicated in both acute and chronic neurodegenerative disorders, including Alzheimer’s disease, Parkinson’s disease and amyotrophic lateral sclerosis (Hynd et al., 2004; Iovino et al., 2020; Van Den Bosch et al., 2006), as well as in retinal diseases (RD) (Boccuni e Fairless, 2022; Telegina, et al. 2022).

Being the retina a direct extension of the CNS, it is not surprising that glutamate and its receptors are expressed on almost every cellular subpopulation of the retina (Boccuni e Fairless, 2022). Retinal glutamate homeostasis is ensured by the concerted action of multiple systems including the counterbalancing action of the inhibitory γ-aminobutyric acid (GABA) neurotransmitter in amacrine cells and some subtypes of bipolar cells, the activity of Müller glia, which accounts for the main scavenging system for the excessive glutamate from the synaptic cleft (Vecino et al., 2016; Boccuni e Fairless, 2022). Moreover, glutamate can *per se* modulate its own release through the activation of NMDARs. Indeed, NMDARs are expressed both on the presynaptic and the postsynaptic neuron, with the latter being involved in signal transduction and the former being autoreceptors that enable the self-adjustment of glutamate release into the cleft (Kesner et al., 2020; Banerjee et al., 2016).

Multiple evidence coming from different animal models of RDs has shown that glutamate imbalance and excitotoxicity are involved in the pathogenesis of retinal diseases (RDs). Intravitreal NMDA injections recapitulates glaucoma in rodents (Mohamad et al., 2021; McKinnon et al., 2009; Tsai et al., 2024). Rat models of geographic atrophy (GA) show increased glutaminase and decreased glutamine synthetase levels (Telegina et al., 2022). Elevated glutamate levels have been noted in diabetic *db/db* mice and streptozotocin-induced diabetic rats (Bogdanov et al., 2014; Ng et al., 2004). Retinal ischemia models revealed reduced glutamate uptake and glutamine synthetase activity (Fernandez et al. 2009), while exogenous glutamate administration has been shown to mimic ischemia (Dorfman et al., 2013). A glutamate receptor antagonist was also shown to prevent visual circuit remodeling and photoreceptor degeneration in the retina of *rd10* mice, which are a widespread model of retinitis pigmentosa (RP; Li et al., 2024). The multiple and compelling evidence supporting the role of glutamate in retinal degeneration has paved the way to the entry of NMDAR antagonists into clinical trials for the treatment of RDs. However, to date, none of the commercially available anti-glutamate drugs has passed clinical trials for RDs, including memantine (Weinreb et al., 2018).

We previously established a novel mechanism contributing to glutamate spillover called “non-canonical presynaptic-induced glutamate spillover” (nPING). According to our model, the aberrant stimulation of presynaptic NMDARs causes the activation c-Jun N-terminal Kinase 2 (JNK2) via the scaffolding protein JNK Interacting Protein 1 (JIP1) (Morel et al., 2018; Kennedy et al., 2007). Once activated, JNK2 phosphorylates the synaptic protein Syntaxin-1A (STX1A) on Ser^14^ (Marcelli et al. 2019). This phosphorylation event, in turn, influences the binding of STX1A to other synaptic proteins, including Synaptosomal-Associated Protein, 25kDa (SNAP25) and Synaptobrein-2 (VAMP2), which have both been shown to bind STX1A more efficiently when phosphorylated on Ser^14^ (Dubois et al., 2002; Foster et al., 1998) and facilitates the mobilization of synaptic vesicles from reserve pools to active zones (Shi et al., 2021). Altogether these events lead to self-amplifying cascade, in which the more glutamate accumulates into the synaptic cleft, the more it elicits its own release (Nisticò et al., 2015).

To specifically investigate such a mechanism, we studied via molecular docking analyses the “minimal contact area” between JNK2 and the N-terminal portion of STX1A and we designed a new small peptide sequence, able to disrupt the interaction between these two proteins. Then, the peptide sequence was been linked to the highly-permeable sequence of the Tat protein of HIV-1, resulting in a cell-permeable peptide called “JGRi1” (Marcelli et al., 2019). Our studies showed that JGRi1 was able to disrupt the interaction between JNK2 and STX1A, reduce STX1A phosphorylation and to reduce NMDA-evoked vesicular release in both human cell lines and in murine brain-derived synaptosomes (Marcelli et al., 2019; Cimino e Feligioni, 2024); Moreover, JGRi1 prevented the NMDA-induced glutamate release in murine brain-derived synaptosomes and, when administered intraperitoneally, it was also able to reach the brain of C57BL/6J (Marcelli et al. 2019).

This work aimed at evaluating whether JGRi1, when topically administered as eye drops, can penetrate the eye and reach the retina and whether, targeting the JNK2-STX1A interaction, can grant neuroprotection in two distinct models of retinal degeneration, namely an ex vivo optic nerve cut model (evONC) and an intravitreal NMDA injection model, when administered topically in the form of eye-drops.

## RESULTS

### The *ex vivo* optic nerve cut model reduces RGC viability and increases L-glutamate immunoreactivity in the retina

Previously, work from our group established a simple *ex vivo* model of retinal degeneration, as shown in Fig.1A. Enucleated eyeballs, following a concomitant optic nerve cut (ONC) showed a powerful RGC loss associated with a strong reduction of their specific markers such as Brain-Specific Homeobox/POU Domain Protein 3A (BRN3A) and Neuronal Nuclei Antigen (NeuN), upregulation of pro-apoptotic marker levels, including cleaved Caspase-3 (c-CASP3) (Fig. 1A; Hassanzadeh et al., 2022; Buccarello et al., 2021).

**Figure 1.**
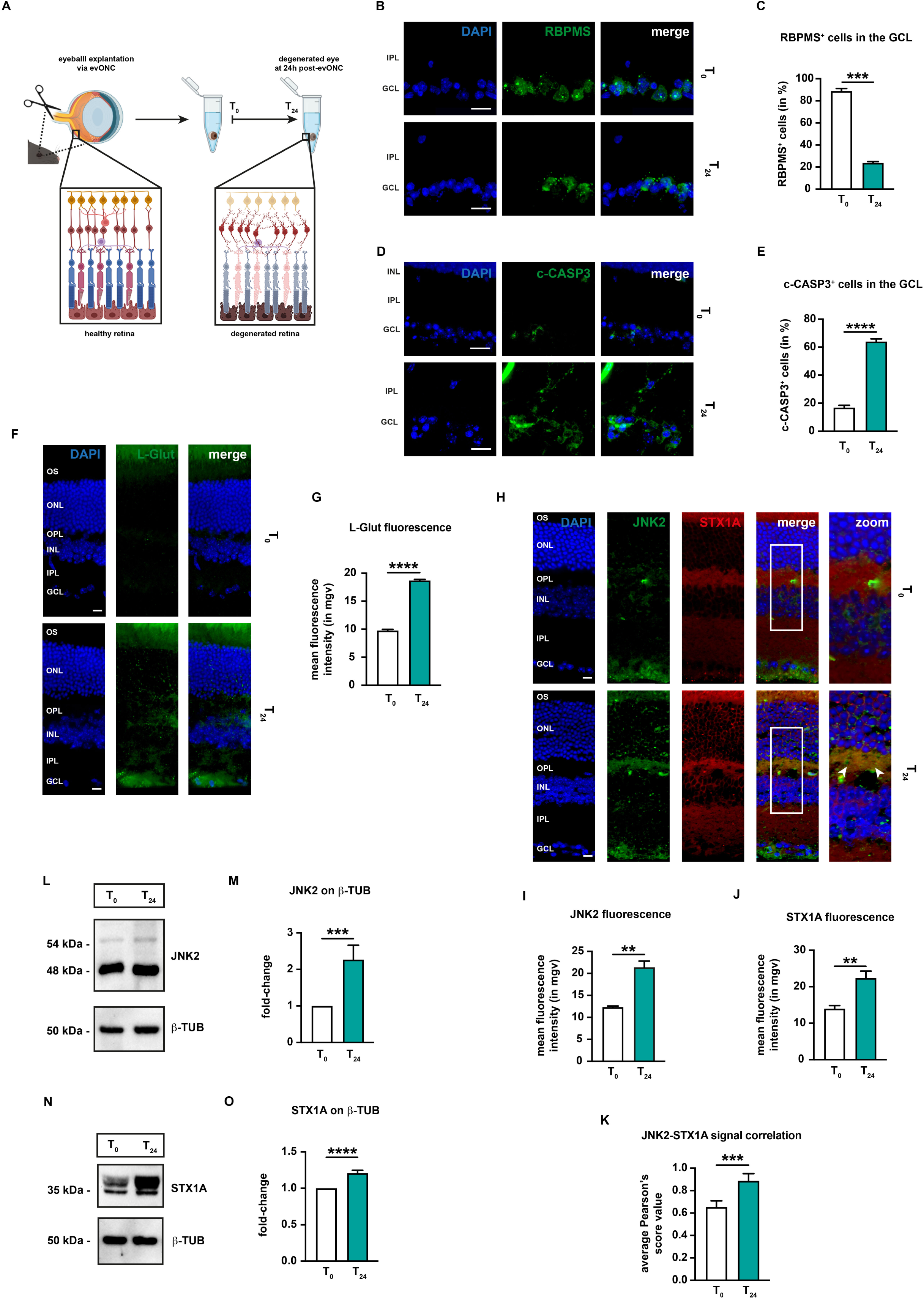
JNK2 and STX1A are induced and co-localize in retinas undergone evONC. **(A)** Schematic representation of the evONC model. **(B)** Representative immunofluorescence for RBPMS. Eyes were processed for immunofluorescence with an RBPMS-specific antibody (green). n = 4 independent experiments. **(C)** Quantification of RBPMS-positive cells. n = 4 independent experiments. **(D)** Representative immunofluorescence for c-CASP3. Eyes were processed for immunofluorescence with a c-CASP3-specific antibody (green). **(E)** Quantification of c-CASP3-positive cells. n = 4 independent experiments. **(F)** Representative immunofluorescence for glutamate. Eyes were processed for immunofluorescence with a glutamate (L-glut) specific antibody (green). n = 4 independent experiments. **(G)** Quantification of the mean fluorescence intensity of L-glut. n = 4 independent experiments. **(H)** Representative co-immunofluorescence for JNK2 and STX1A. Eyes were processed for immunofluorescence with a JNK2-specific antibody (green) and a STX1A-specific antibody (red). White arrowheads in the zoom column indicate areas where JNK2 and STX1A signals co-localize. n = 5 independent experiments. **(I)** Quantification of the mean fluorescence intensity of JNK2. n = 5 independent experiments. **(J)** Quantification of the mean fluorescence intensity of STX1A. n = 5 independent experiments. **(K)** Correlation analysis between JNK2 and STX1A signals. Pearson’s scores were calculated per each image from (I). n = 5 independent experiments. **(L)** Representative western blot for JNK2. Samples were blotted, then incubated with primary antibodies against JNK2 and β-TUB (loading control). n = 4 independent experiments. **(M)** Densitometric analysis of JNK2 with respect to β-TUB. n = 4 independent experiments. **(N)** Representative western for STX1A. Samples were blotted, then incubated primary antibodies against STX1A and β-TUB (loading control). n = 4 independent experiments. **(O)** Densitometric analysis of STX1A with respect to β-TUB. n = 4 independent experiments. **Immunofluorescence caption:** OS, outer segment; ONL, outer nuclear layer; OPL, outer plexiform layer; INL, inner nuclear layer; IPL, inner plexiform layer; GCL, ganglion cell layer. 40X magnification. Scale bar 10 μM. **Bar plot caption:** Bar representing mean +/− S.D. Statistical analysis: unpaired t-test, p < 0.05.

Retinal tissue sections are frequently subjected to autofluorescence, primarily deriving from the fact that the retina is rich in autofluorescent species, such as lipofuscin and melanin (Petty et al., 2011). Therefore, to ascertain the reliability of our immunofluorescence data, immunofluorescence analysis was carried out on retinal slices without any antibody staining. Our analysis showed that retinal autofluorescence was mainly localized within the outer segment (OS) of photoreceptors (Fig. S1A). Therefore, fluorescent signal in the OS will not be considered within this work. Additionally, to exclude possible artifacts deriving from secondary antibody cross-reactivity, retinal slices were incubated only with secondary antibodies. As shown in fig. S1B, no antibody cross-reactivity was detectable within our samples.

In line with previous work, at 24 hours post-ONC (T_24_), we found that RGC viability was reduced in comparison to non-degenerated eyes (T_0_). Indeed, the number of cells positive to the RGC marker RNA-Binding Protein with Multiple Splicing (RBPMS) in the ganglion cell layer (GCL) were reduced (Fig. 1B-C). Whereas, the c-CASP3-positive cells in the GCL concomitantly increased (Fig.1D-E). All this evidence combined confirmed that our evONC model fosters RGC degeneration, in line with our previous reports (Hassanzadeh et al., 2022; Buccarello et al., 2021).

Afterwards, given the involvement of glutamate unbalance in the pathogenesis of several RD (Boccuni e Fairless, 2022; Ishikawa, 2013), we investigated potential changes in glutamate immunoreactivity in retina upon ONC. The retinal glutamate level was measured by using a specific L-glutamate antibody in immunofluorescent experiments. Our analysis showed that at T_0_ glutamate signal poorly detectable, while at T_24_ glutamate appeared as a more punctate staining present throughout the entire tissue, accumulating especially at the level of the GCL and the outer plexiform layer (OPL) (Fig.1F). Quantification of the mean fluorescence intensity of L-glutamate validated our initial findings (Fig. 1G), confirming that upon evONC there in an increase in L-glutamate immunoreactivity.

### JNK2 and STX1A are upregulated in the *ex vivo* ONC mouse model

Afterwards, the expressions of both JNK2 and STX1A was evaluated in evONC retinas via co-immunofluorescence at both T_0_ and T_24_. As shown in Fig. 1H, JNK2 signal was present mainly in the GCL at T_0_, although some signal was detectable from the inner nuclear layer (INL) and outer plexiform layer (OPL); STX1A, on the other hand, was expressed throughout the retina and it was particularly abundant in the inner plexiform layer (IPL) and OPL. At T_24_, JNK2 signal strongly increased in the OPL, while STX1A immunoreactivity appeared higher through all the layers of the retina (Fig.1I), including the OPL, where it exhibited a striking co-localization with JNK2 (Fig.1H - merge, white arrowheads). The change in JNK2 and STX1A signals was also assessed by measuring the mean fluorescence intensity of both signals. Our analysis confirmed the increase of the fluorescence intensity of both JNK2 (Fig. 1I) and STX1A (Fig. 1J). Finally, the co-localization between the JNK2 and STX1A signals was assessed by calculating the Pearson’s score. As expected, at T_24_, there was an increase in the average Pearson’s score value in our slices (Fig. 1K).

Furthermore, the retinal expression of JNK2 and STX1A upon evONC was also assessed by immunoblot. JNK2 at T_0_ appeared as two distinct bands, in line with previous reports (Kallunki et al., 1994), one with higher intensity with a molecular weight of around 46 kDa and second one at around 54 kDa. At T_24_, there was an overall increase in the band signal intensity, which proved to be statistically significant upon densitometric analysis (Fig. 1L-M). On the other hand, STX1A appeared as a two very close bands with molecular weight of around 35 kDa. The intensity of these bands increased upon ONC (Fig.1N-O).

### NMDA treatment fosters RGC degeneration and glutamine synthetase expression in retinal wholemount preparations

The contribution of NMDA to retinal degeneration was assessed using cultured retinal wholemount preparations undergone a 2-hour treatment with 100 μM NMDA. After 24 hours, retinas were processed accordingly (Fig. 2A-B). Firstly, RGC viability was assessed by immunoblot for the RGC marker Brain-Specific Homeobox/POU Domain Protein 3A (BRN3A). The analysis revealed that at 24 hours post-mounting there was a dramatic reduction of BRN3A expression, which was enhanced by treatment with 100 μM NMDA (Fig. 2C-D).

**Figure 2.**
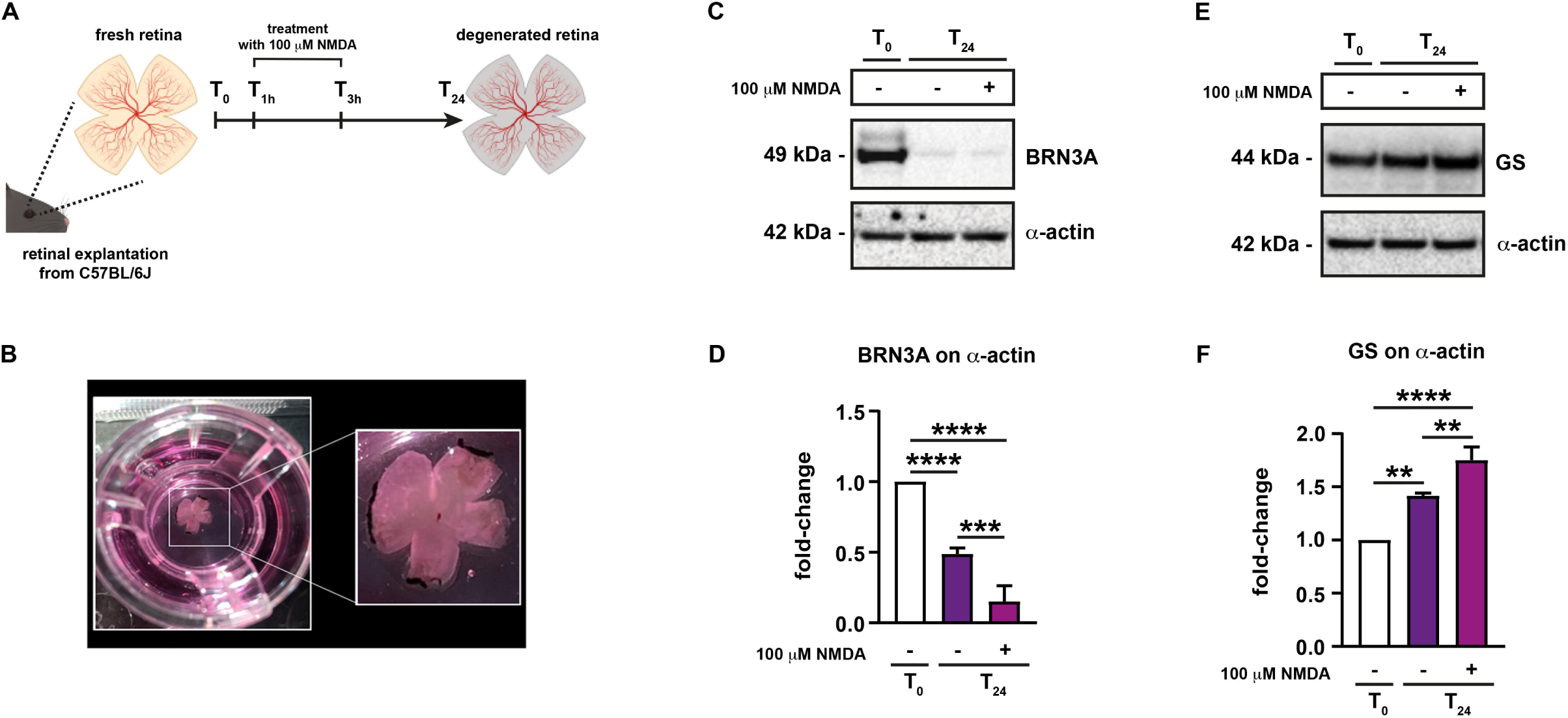
NMDA treatment induces RGC degeneration and GS expression in retina wholemount preparations. **(A)** Schematic representation of cultured retina wholemount preparations. **(B**) Representative picture of a cultured retinal wholemount preparation. **(C)** Representative western blot for BRN3A and GS. Samples were blotted, then incubated with a specific primary antibody against BRN3A, GS and β-TUB (loading control). n = 5 independent experiments. **(D)** Densitometric analysis of BRN3A with respect to β-TUB. n = 5 independent experiments. **(E)** Densitometric analysis of GS with respect to β-TUB. Bar plots representing mean +/− S.D. n = 4 independent experiments. **Bar plot caption:** Bar representing mean +/− S.D. Statistical analysis: One-way ANOVA, *post hoc* Tukey test, p < 0.05.

Additionally, the expression of glutamine synthetase (GS), one of the main enzymes involved in converting glutamate to glutamine (Bringmann et al., 2013) was assessed by immunoblot. The analysis revealed that at T_24_ there was a slight upregulation of GS expression, which was more marked by the addition of 100 μM NMDA (Fig. 2E-F).

### Intravitreal injection of NMDA induced retinal neurodegeneration

Another well-established model for retinal degeneration is the NMDA-induced degeneration model which is known to promote RGC degeneration through NMDA overactivation and glutamate overflow (Fig. 3A). To test retinal neurodegeneration in this model both RGC viability and apoptosis were assessed by using antibodies against specific markers, NeuN and c-CASP3 respectively. 30 days after NMDA injection, there was a reduction in RCG viability in comparison to untreated mice, as shown by both immunofluorescence for the RGC marker NeuN and confirmed by the quantification of NeuN-positive cells in the GCL (Fig.3B-C); conversely, at 30 days after the NMDA injection, the number of c-CASP3-positive cells in the GCL was higher than untreated mice (Fig. 3D-E). To address the effect of NMDA injection on neuronal connections within the retina, 4 days prior to their sacrifice, mice were injected with fluorescently labeled cholera toxin B (CTB-488), which enabled anterograde tracing of RGC axons in the retinal nerve fiber layer (RFNL). In untreated retinas, CTB-488 was taken up by RGCs and effectively transported through their axons in the RFNL, whilst in NMDA-injected mice CTB-488 uptake and transport failed in RGCs, indicating RGC compromise following NMDA exposure (Fig.3F; Saggu et al., 2010), which was also confirmed by the subsequent quantification of CTB-488 fluorescence (Fig.3G).

**Figure 3.**
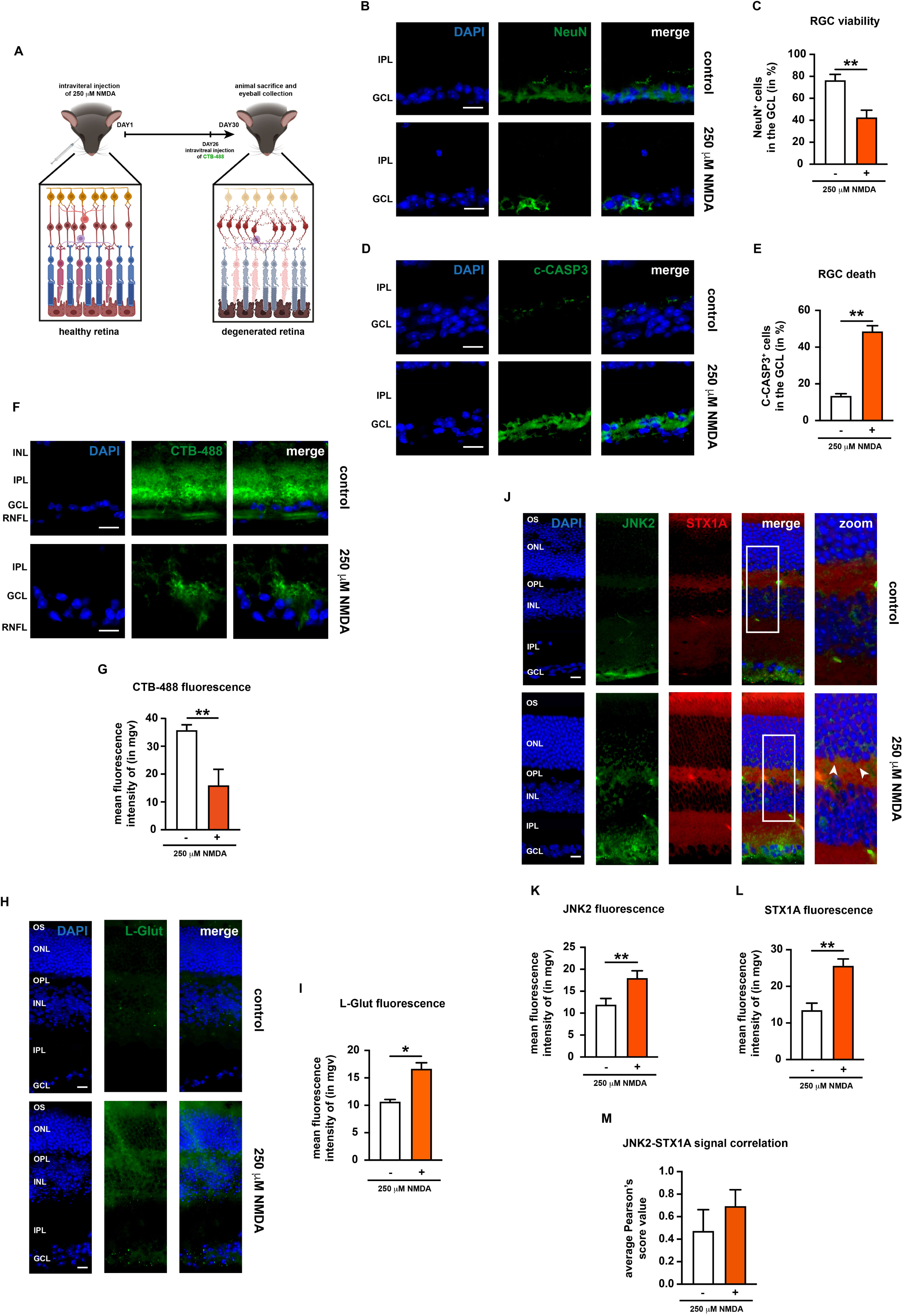
NMDA injection fosters retinal degeneration and induces JNK2 and STX1A expression. **(A)** Schematic representation of the NMDA injection model. **(B)** Representative immunofluorescence for NeuN. Eyes were processed for immunofluorescence with a NeuN-specific antibody (green). n = 4 independent experiments. 40X magnification. **(C)** Quantification of NeuN-positive cells. n = 4 independent experiments. **(D)** Representative immunofluorescence for c-CASP3. Eyes were processed for immunofluorescence with a c-CASP3-specific antibody (green). n = 4 independent experiments. **(E)** Quantification of c-CASP3-positive cells. n = 4 independent experiments. **(F)** Representative immunofluorescence for CTB-488. n = 5 independent experiments. **(G)** Quantification of the mean fluorescence intensity of CTB-488. n = 5 independent experiments. **(H)** Representative immunofluorescence for glutamate. Eyes were processed for immunofluorescence with a glutamate (L-glut)-specific antibody (green). n = 4 independent experiments. **(I)** Quantification of the mean fluorescence intensity of L-glut. n = 4 independent experiments. **(J)** Representative co-immunofluorescence for JNK2 and STX1A. Eyes were processed for immunofluorescence with a JNK2 specific antibody (green) and a STX1A specific antibody (red). White arrowheads in the zoom column indicate areas where JNK2 and STX1A signals strongly co-localize. n = 5 independent experiments. **(K)** Quantification of the mean fluorescence intensity of JNK2. n = 5 independent experiments. **(L)** Quantification of the mean fluorescence intensity of STX1A. n = 5 independent experiments. **(M)** Correlation analysis between JNK2 and STX1A signals. Pearson’s scores were calculated per each image from (J). n = 5 independent experiments. **Immunofluorescence caption:** OS, outer segment; ONL, outer nuclear layer; OPL, outer plexiform layer; INL, inner nuclear layer; IPL, inner plexiform layer; GCL, ganglion cell layer. 40X magnification. Scale bar 10 μM. **Bar plot caption:** Bar representing mean +/− S.D. Statistical analysis: unpaired t-test, p < 0.05.

To assess the effect of NMDA on retinal glutamate levels, immunofluorescence for L-glutamate was carried out. In untreated retinas, L-glutamate was poorly detectable, although some signal was detectable in the inner retina, between the INL and the OPL; however, in NMDA-injected mice, there was a dramatic increase in L-glutamate immunoreactivity, which was identifiable not only in the inner retina, but in the GCL as well, although to a lesser extent (Fig. 3H). Such data were confirmed by the quantification of the mean fluorescence intensity of L-glutamate signal in retinal samples (Fig. 3I).

### Intravitreal injection of NMDA promotes the expression of JNK2 and STX1A in the retina

The expression of JNK2 and STX1A were assessed in the retina of untreated and NMDA-injected mice. In untreated mice, JNK2 expression was detectable mainly in the GCL, while after intravitreal NMDA injection JNK2 immunoreactivity engaged the IPL and OPL, although a few JNK2 signal was detectable even in the INL and ONL as well (Fig. 3J). Such NMDA-dependent increase in JNK2 immunoreactivity was also confirmed by quantification of mean fluorescence intensity in the retina (Fig. 3K). Similarly to JNK2, STX1A immunoreactivity was also higher in the retinas of NMDA-injected mice. Like for JNK2, quantification of STX1A fluorescence in the retina confirmed the NMDA-induced increase of STX1A immunoreactivity (Fig. 3L). Afterwards, the JNK2-STX1A co-localization in the retina of NMDA-injected mice was analyzed via by Pearson’s analysis, which showed that upon NMDA injection there was a slight, yet no significant, increase in the co-localization between the two proteins (Fig. 3M).

### JGRi1 effectively reaches the retina upon *ex vivo* and *in vivo* administration

In previous works, we have showed that the cell-permeable peptide JGRi1 was effective in preventing the JNK2-STX1A interaction and, therefore, counteracting glutamate spillover in multiple models (Marcelli et al., 2019; Cimino e Feligioni, 2024). On these premises and being both the JNK2-STX1A interaction and glutamate release induced in the retina of mouse model used in this work, we decided to test the potential neuroprotective effect of JGRi1 in our experiments.

Firstly, the eye permeability of JGRi1 has been tested in C57BL/6J mice upon *ex vivo* treatment. Enucleated mouse eyeballs were immersed into balanced salt solution (BSS) buffer and incubated with different dosages (50 μM, 250 μM, 500 μM) of fluorescein (FITC)-tagged JGRi1 (F-JGRi1) for 1 hour at 37°C (Fig. 4A). As a control, we included in our experimental design a version of the peptide devoid of the HIV-Tat sequence (ΔTat-F-JGRi1), with consequent reduced cell permeability. Immunofluorescence analysis carried out on retinal slices revealed that FITC-JGRi1 was able to reach the retina upon topical application. JGRi1 was detected in multiple layers, including GCL, the IPL and the OPL (Fig. 4B). Conversely, incubation of enucleated eyeballs with ΔTat-F-JGRi1 resulted in very low accumulation of F-JGRi1 signal in the retina (Fig. 4B). The amount of peptide that reached the retina upon treatment was quantified: F-JGRi1 accumulated in the both the OPL and the GCL in a dose-dependent manner, whereas, as expected, ΔTat-F-JGRi1 could not (Fig. 4C-D). Then, the number of FITC-positive cells in the GCL was measured in the immunofluorescence experiments (Fig. 4E) showing that the treatment with increasing dosages of F-JGRi1 led to a dose-dependent increase of FITC-positive cells in the GCL (Fig. 4F) while, as expected, treatment with ΔTat-F-JGRi1 resulted in almost no accumulation of the peptide within the GCL (Fig. 4F).

**Figure 4.**
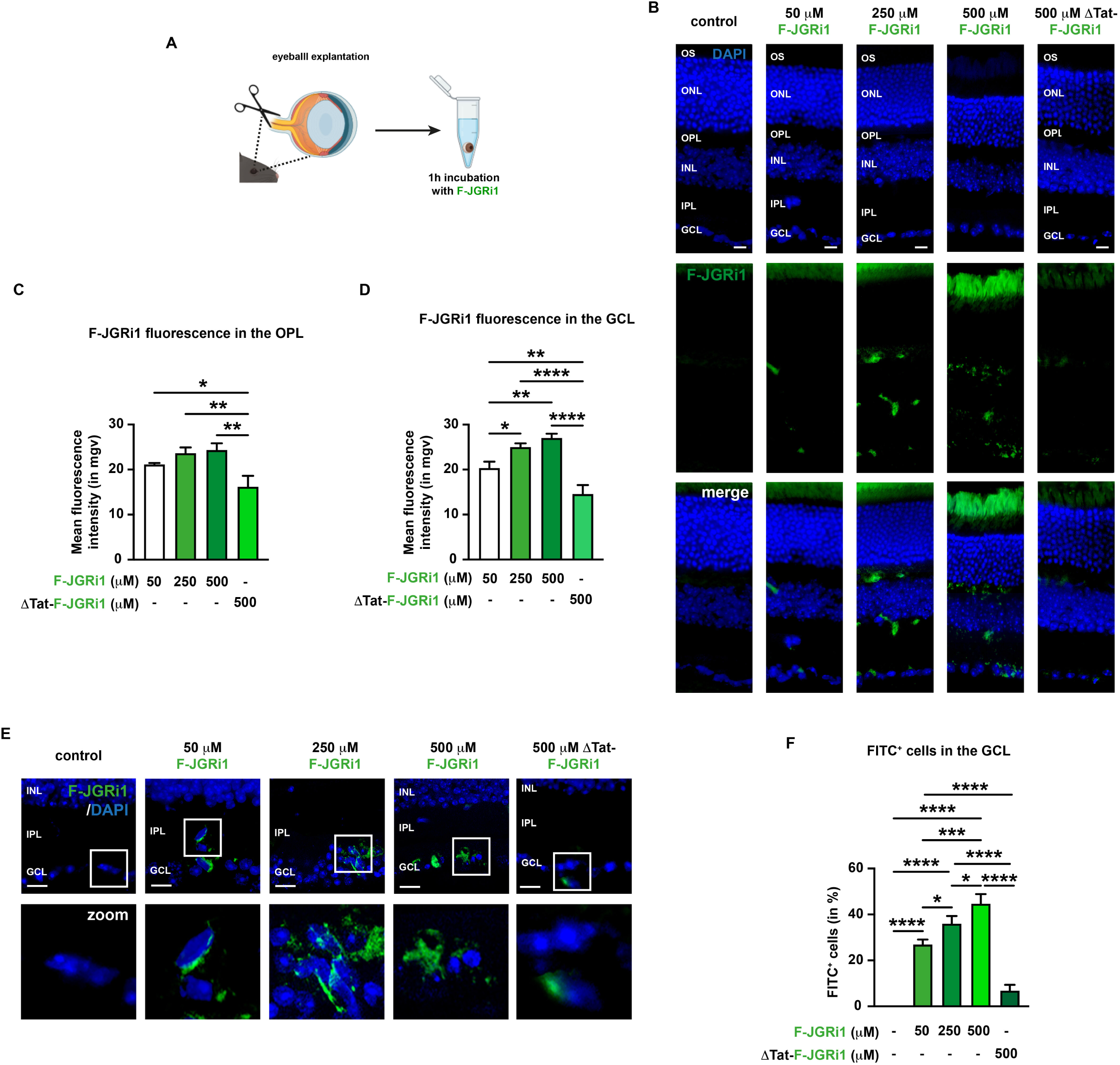
Fluorescently-labelled JGRi1 (F-JGRi1) penetrates the eye and accumulates in the retina of C57BL/6J mice upon *ex vivo* treatment. **(A)** Schematic representation of F-JGRi1 treatment. **(B)** Representative immunofluorescence for F-JGRi1. n = 5 independent experiments. **(C)** Quantification of the mean fluorescence intensity of F-JGRi1 in the OPL. n = 5 independent experiments. **(D)** Quantification of the mean fluorescence intensity of F-JGRi1 in the GCL. n = 5 independent experiments. (**E)** Enlarged view of F-JGRi1 accumulation in the GCL. n = 5 independent experiments. **(F)** Quantification of F-JGRi1-positive cells. n = 5 independent experiments. **Immunofluorescence caption:** OS, outer segment; ONL, outer nuclear layer; OPL, outer plexiform layer; INL, inner nuclear layer; IPL, inner plexiform layer; GCL, ganglion cell layer. 40X magnification. Scale bar 10 μM. **Bar plot caption:** Bar representing mean +/− S.D. Statistical analysis: One-way ANOVA, *post hoc* Tukey test, p < 0.05.

Afterwards, the murine eye permeability of JGRi1 was also tested upon topical administration in *in vivo* treatment to determine its accumulation in the retina. For this purpose, C57BL/6J mice were treated with different dosages of FITC-JGRi1 (50 μM, 250 μM, 500 μM) dissolved into BSS buffer in the form of eye-drops for 6 days, one drop *per die*. Afterwards, animals were sacrificed and the obtained eyeballs were analyzed by immunofluorescence. ΔTat-F-JGRi1 was included as a control (Fig. 5A). Similarly to what was observed in the *ex vivo* treatment experiment, topically administered F-JGRi1 successfully accumulated in the retina, especially in the GCL and in the OPL (Fig. 5B). Of note, differently from the ex vivo treatment, some ΔTat-F-JGRi1 accumulated in the GCL, but not in the OPL upon treatment (Fig. 5B), suggesting that, even without the Tat sequence, JGRi1 has some cell-permeability properties upon prolonged treatment. Quantification of the FITC signal throughout the retina showed a slight dose-dependent accumulation of F-JGRi1 in the OPL and the GCL (Fig. 5C-D). Similarly to what was done for the *ex vivo* treatment, the FITC-positive cells were counted in the GCL upon JGRi1 treatment. As outcome, the increase of FITC-positive cells in the GCL, in line with our previous *ex vivo* data, follows a dose-dependent trend (Fig. 3F-G).

**Figure 5.**
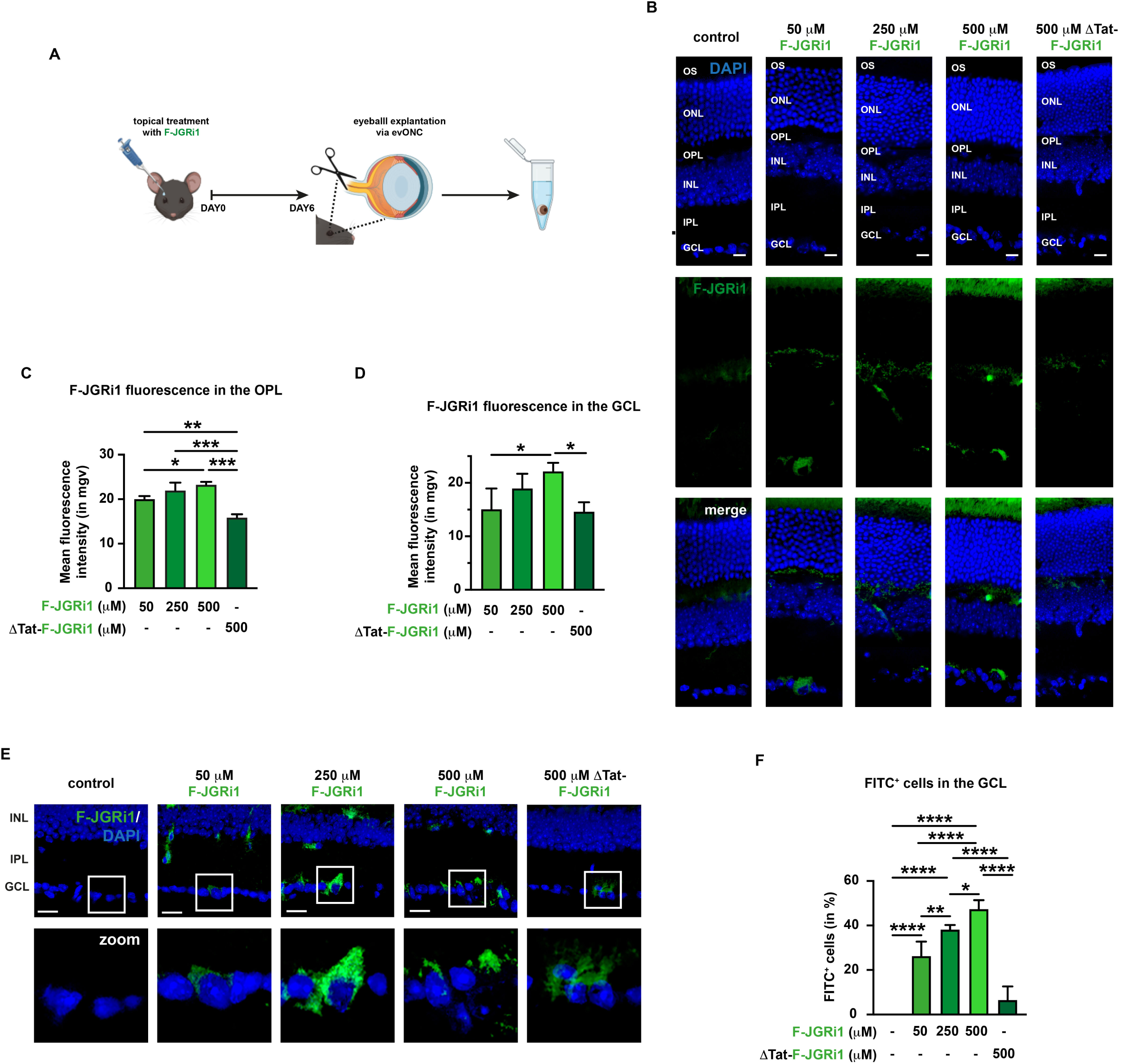
F-JGRi1 penetrates the eye and accumulates in the retina of C57BL/6J mice upon topical administration for 6 days. **(A)** Schematic representation of in vivo F-JGRi1 treatment. **(B)** Representative immunofluorescence for F-JGRi1. n = 5 independent experiments. **(C)** Quantification of the mean fluorescence intensity of F-JGRi1 in the OPL. n = 5 independent experiments. **(D)** Quantification of the mean fluorescence intensity of F-JGRi1 in the GCL. n = 5 independent experiments. (**E)** Enlarged view of F-JGRi1 accumulation in the GCL of treated mice. n = 5 independent experiments. **(F)** Quantification of F-JGRi1-positive cells. n = 5 independent experiments. **Immunofluorescence caption:** OS, outer segment; ONL, outer nuclear layer; OPL, outer plexiform layer; INL, inner nuclear layer; IPL, inner plexiform layer; GCL, ganglion cell layer. 40X magnification. Scale bar 10 μM. **Bar plot caption:** Bar representing mean +/− S.D. Statistical analysis: One-way ANOVA, *post hoc* Tukey test, p < 0.05.

### Topical treatment with JGRi1 protects RGC from evONC-induced degeneration

Next, the protective effect of JGRi1 was assessed in the evONC-induced retinal degeneration mouse model. C57BL/6J mice were treated one drop daily for 6 days with JGRi1 at different concentrations (50 μM, 250 μM and 500 μM). Afterwards, animals were sacrificed and eye bulbs, removed by evONC, were incubated overnight at 4°C in PBS to allow their degeneration. A scrambled version of the JGRi1 peptide (sJGRi1) was also included as control (Fig. 6A).

**Figure 6.**
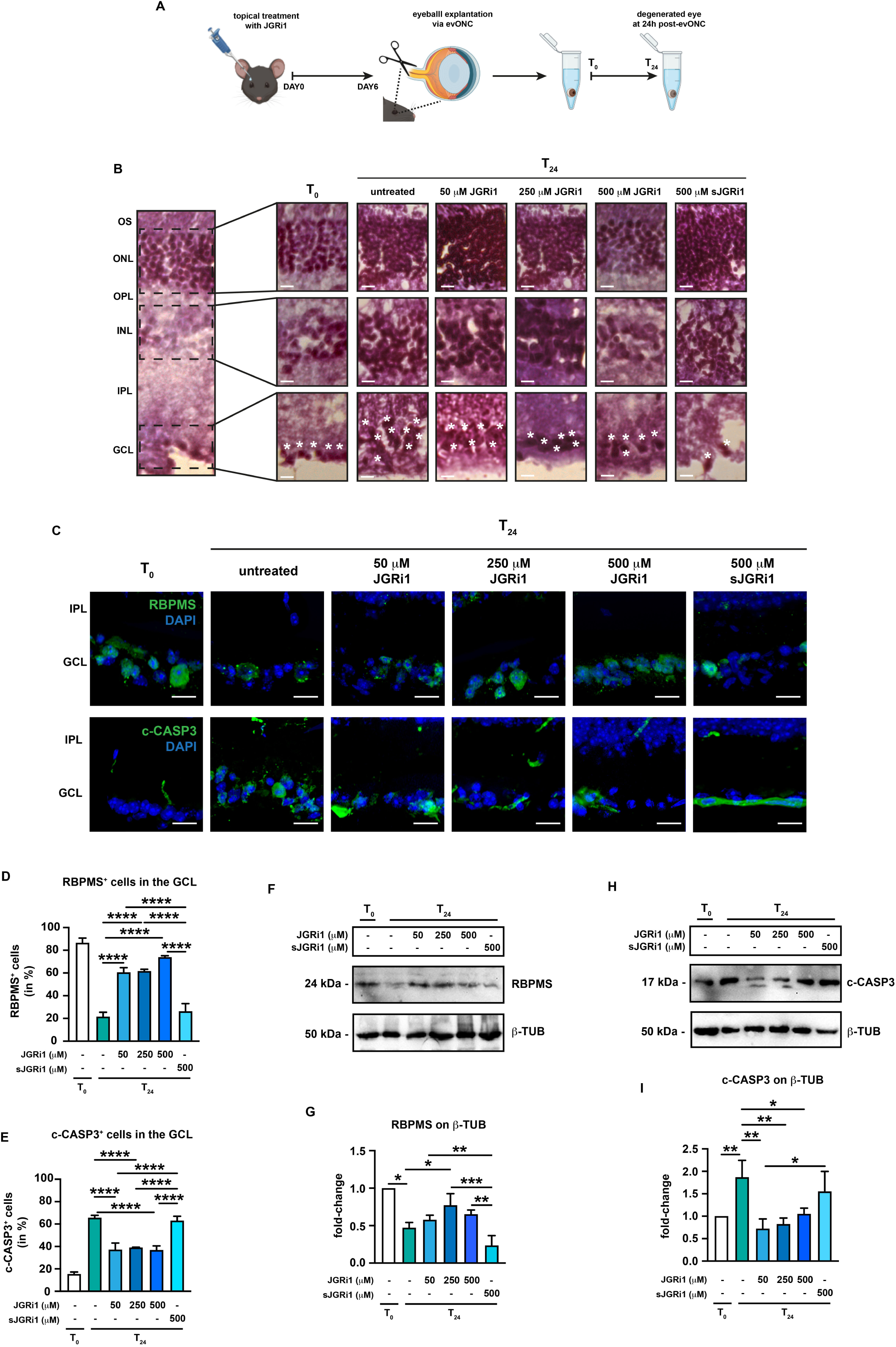
JGRi1 protects retinas and RGCs from evONC-induced neurodegeneration. **(A)** Schematic representation of JGRi1 treatment. **(B)** Representative hematoxylin-eosin staining. Eyes were stained with hematoxylin, counterstained with eosin. A sample retina is shown in the left part of the panel. For each nuclear layer an enlarged view is provided. White asterisks indicate RGCs. n = 3 independent experiments. **(C)** Representative immunofluorescence for (upper row) and c-CASP3 (bottom row) on retinal slices from evONC model. Eyes were processed for immunofluorescence with specific antibody (green). n = 4 independent experiments. **(D)** Quantification of RBPMS-positive cells. n = 4 independent experiments. For reader’s convenience statistical significance only in comparison to T_24_ and between active JGRi1 and sJGRi1 is shown. **(E)** Quantification of c-CASP3-positive cells. n = 4 independent experiments. For reader’s convenience statistical significance only in comparison to T_24_ and between active JGRi1 and sJGRi1 is shown. **(F)** Representative western blot for RBPMS. Samples were blotted, then incubated with primary antibodies against RBPMS and β-TUB (loading control). n = 4 independent experiments. **(G)** Densitometric analysis of RBPMS with respect to β-TUB. n = 4 independent experiments. **(H)** Representative western blot showing the expression of c-CASP3 in retinas from evONC model. Samples were blotted, then incubated with primary antibodies against c-CASP3 and β-TUB (loading control). n = 5 independent experiments. **(G)** Densitometric analysis of c-CASP3 with respect to β-TUB. n = 5 independent experiments. **Immunofluorescence caption:** OS, outer segment; ONL, outer nuclear layer; OPL, outer plexiform layer; INL, inner nuclear layer; IPL, inner plexiform layer; GCL, ganglion cell layer. 40X magnification. Scale bar 10 μM. **Bar plot caption:** Bar representing mean +/− S.D. Statistical analysis: One-way ANOVA, *post hoc* Tukey test, p < 0.05.

Firstly, hematoxylin/Eosin staining was carried out to evaluate the retinal cytoarchitecture in the experimental conditions. The results showed that at T_24_ there was a loss of integrity within all three nuclear layers of the retina while the treatment with increasing dosages of JGRi1 led to a progressive restoration of the tissue architecture of the retinal tissue (Fig. 6B).

Immunofluorescence for RBPMS was carried out to evaluate RGC viability. As already shown in fig. 1C, also this set of experiments demonstrated that at T_24_ there was RGC loss, visualized as the reduction of RBPMS-positive cells in the GCL in comparison to T_0_. Interestingly, treatment with all three doses of JGRi1 caused an increase of the number RBPMS-positive cells in the GCL, with no significant differences between all the tested doses. As expected, treatment with 500 μM sJGRi1 failed to rescue RGC viability upon evONC (Fig. 6C-D). RBPMS expression was also evaluated by immunoblot in which RBPMS appeared as a single band at around 24 kDa of molecular weight. The densitometric analyses showed that RBPMS was reduced at T_24_, while JGRi1 led to a recovery of RBPMS signal, although only 250 μM JGRi1 provided statistical significance (Fig. 6F-G). As expected, the treatment with sJGRi1 did not produce any protective effect on RGC viability in comparison to the active peptide. Meanwhile, the number of apoptotic cells in the GCL was addressed by measuring the immunofluorescence for c-CASP3. As expected, at T_24_, the number of c-CASP3-positive was higher, and treatment with all dosages of JGRi1 was able to reduced them in a significant manner (Fig. 6C-E). In parallel, c-CASP3 expression was evaluated by immunoblotting using an antibody specific for the cleaved form of the protein, which yielded a single band with a molecular weight of 17 kDa. c-CASP3 expression increased at T_24_, while it was reduced significantly by all three doses of JGRi1, in line with the immunofluorescence reports. Even in this case, the treatment with sJGRi1 did not produce any effect on the evONC-induced c-CASP3 upregulation.

### Topical JGRi1 reduces the evONC-induced glutamate release

Afterwards, the effect of JGRi1 on the evONC-induced release of glutamate was addressed by immunofluorescence. At T_24_, as expected, glutamate immunoreactivity was higher throughout the entire tissue (Fig. 7A), but treatment with JGRi1 progressively reduced it. Of note, only 250 μM and 500 μM JGRi1 produced a significative reduction in retinal glutamate immunoreactivity, with the latter bringing it even below control levels (Fig. 7B).

**Figure 7.**
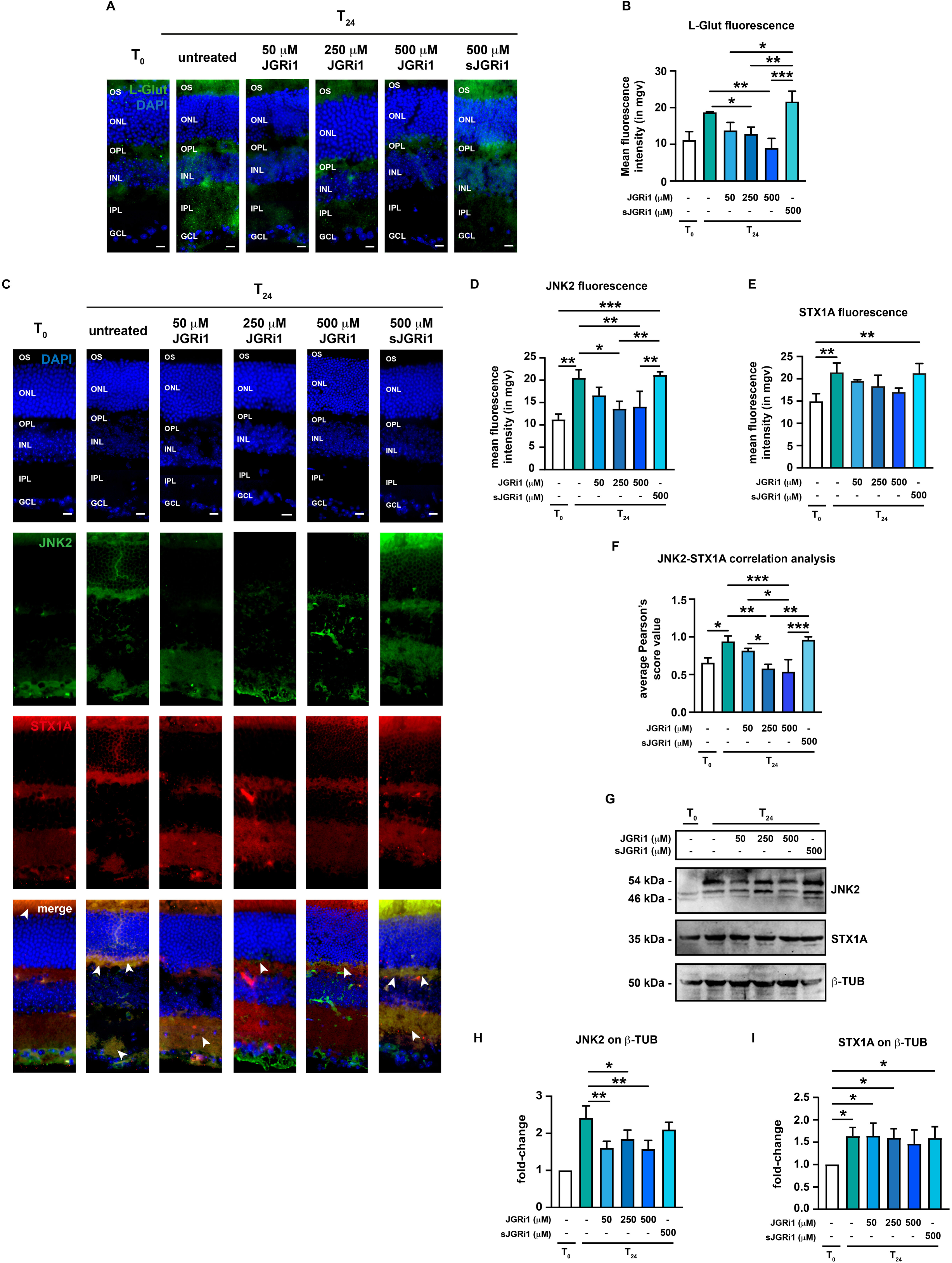
Effect of JGRi1 on retinal glutamate levels and the expression JNK2 and STX1A in evONC model. **(A)** Representative immunofluorescence for L-glut. Eyes were processed for immunofluorescence with an L-glut-specific antibody (green). n = 3 independent experiments. **(B)** Quantification of the mean fluorescence intensity of L-glut. n = 3 independent experiments. **(C)** Representative co-immunofluorescence for JNK2 and STX1A on retinal slices from evONC model. Eyes were processed for immunofluorescence with a JNK2 specific antibody (green) and a STX1A specific antibody (red). White arrowheads in the merge column indicate areas where JNK2 and STX1A signals co-localize. n = 5 independent experiments. **(D)** Quantification of the mean fluorescence intensity of JNK2. n = 5 independent experiments. **(E)** Quantification of the mean fluorescence intensity of STX1A. n = 5 independent experiments. **(F)** Correlation analysis between JNK2 and STX1A signals in retinal slices from evONC. Pearson’s scores were calculated per each image from (C). n = 5 independent experiments. **(G)** Representative western blot showing the expression of JNK2 and STX1A in retinas from evONC model. Samples were blotted, then incubated with primary antibodies against JNK2, STX1A and β-TUB (loading control). n = 5 independent experiments. **(H)** Densitometric analysis of JNK2 with respect to β-TUB. n = 5 independent experiments. For reader’s convenience statistical significance only in comparison to T_24_ and between active JGRi1 and sJGRi1 is shown. **(I)** Densitometric analysis of STX1A with respect to β-TUB. n = 5 independent experiments. **Immunofluorescence caption:** OS, outer segment; ONL, outer nuclear layer; OPL, outer plexiform layer; INL, inner nuclear layer; IPL, inner plexiform layer; GCL, ganglion cell layer. 40X magnification. Scale bar 10 μM. **Bar plot caption:** Bar representing mean +/− S.D. Statistical analysis: One-way ANOVA, *post hoc* Tukey test, p < 0.05.

### Topical administration of JGRi1 reduces the upregulation of JNK2 but not STX1A, as well as their retinal co-localization induced by evONC

To evaluate the effect of JGRi1 on JNK2 and STX1A expression in the retina, evONC eyes were first treated with the afore-mentioned doses of JGRi1 and sJGRi1, then processed for co-immunofluorescence using JNK2- and STX1A-specific antibodies. As shown previously, at T_24_ JNK2 immunoreactivity in the inner retina increased. However, treatment with JGRi1 led to a decrease of JNK2 immunoreactivity throughout the tissue: indeed, while 50 μM JGRi1 only produced a mild reduction in JNK2 immunoreactivity, 250 μM JGRi1 restored a pattern of JNK2 expression very close to that seen at T_0_, although some JNK2 signal remained detectable in the inner retina. Ultimately, treatment with 500 μM JGRi1 reduced JNK2 immunoreactivity in the GCL, although dome signal was still detectable between the OPL and the INL (Fig. 7C). Likewise, STX1A expression increased at T_24_, but, unlike JNK2, a slight yet no significant reduction in STX1A level was found in response to all the tested dosages of JGRi1 (Fig. 7C). Such observations were also validated by measuring the fluorescence intensity of both proteins on retinal sections (Fig. 7D-E). As expected, pre-treatment with sJGRi1 di not have any effect on JNK2 or STX1A expression at T_24_.

As previously shown in fig.1K, JNK2 and STX1A signals strongly co-localized in the OPL at T_24_. Therefore, we decided to address the effect of JGRi1 on the evONC-fostered co-localization between the two proteins. Co-immunofluorescence for JNK2 and STX1A showed that JGRi1 led to a progressive reduction of the co-localization between the two signals in a dose-dependent fashion. In fact, while at 50 μM JGRi1 JNK2 and STX1A still co-localized in the IPL, treatment with 250 μM and 500 μM JGRi1 produced a minimal overlap between their fluorescence signals (Fig. 7C – white arrowheads). Interestingly, upon treatment with sJGRi1, there was an even more marked co-localization between JNK2 and STX1A in comparions to T_0_ (Fig. 7C). A Pearson’s analysis showed that, while the average Pearson’s score increased at T_24_, it diminished in a significant manner upon treatments with 250 μM and 500 μM JGRi1 (Fig.7F).

Afterwards, protein levels of JNK2 and STX1A were both assessed by immunoblot. JNK2 expression, which already showed an increase at T_24_, was reduced by at all dosages of JGRi1, but not by sJGRi1 (Fig. 7G). However, its expression was still higher than the T_0_, probably as a result of the fact that the p54 band was still highly expressed in any condition (Fig. 7G-H). Similarly to JNK2, immunoblot for STX1A also confirmed our immunofluorescence reports, with none of the tested JGRi1 doses able to bring back STX1A expression back to control levels (Fig. 7G-I).

### Topical administration of JGRi1 reduces the phosphorylation of both JNK2 and STX1A and the rate SNARE complex formation induced by evONC

In our previous works, we reported that in the evONC mouse model the phosphorylation of JNK was increased within the retina (Hassanzadeh et al., 2022; Buccarello et al., 2021). Therefore, the effect of JGRi1 treatment of the evONC-induced JNK phosphorylation was assessed by immunoblot. At T_24_ the phosphorylation of JNK was strongly induced in comparison to T_0_; the treatments with all doses of JGRi1, but not sJGRi1, reduced the JNK phosphorylation with significancy reached only with 500 μM JGRi1 (Fig. 8A-B). Notably, treatment with 500 μM JGRi1 seemed to induce a significant upregulation of total JNK levels in the immunoblot, as confirmed by band densitometry (Fig.8A-C). Afterwards, the changes in STX1A phosphorylation at Ser^14^ was also investigated. As said previously, such residue is specifically phosphorylated by JNK2 during excitotoxic stress, thus favoring the release of glutamate (Marcelli et al., 2019; Cimino e Feligioni, 2024). In the immunoblots phosphorylated STX1A (p-STX1A) appeared as a two distinct band of around 35 kDa (Fig. 8D); at T_24_ p-STX1A signal was higher than T_0_, as confirmed by densitometric analysis; p-STX1A levels were lowered only by treatments with 250 μM JGRi1 and 500 JGRi1 (Fig. 8E).

**Figure 8.**
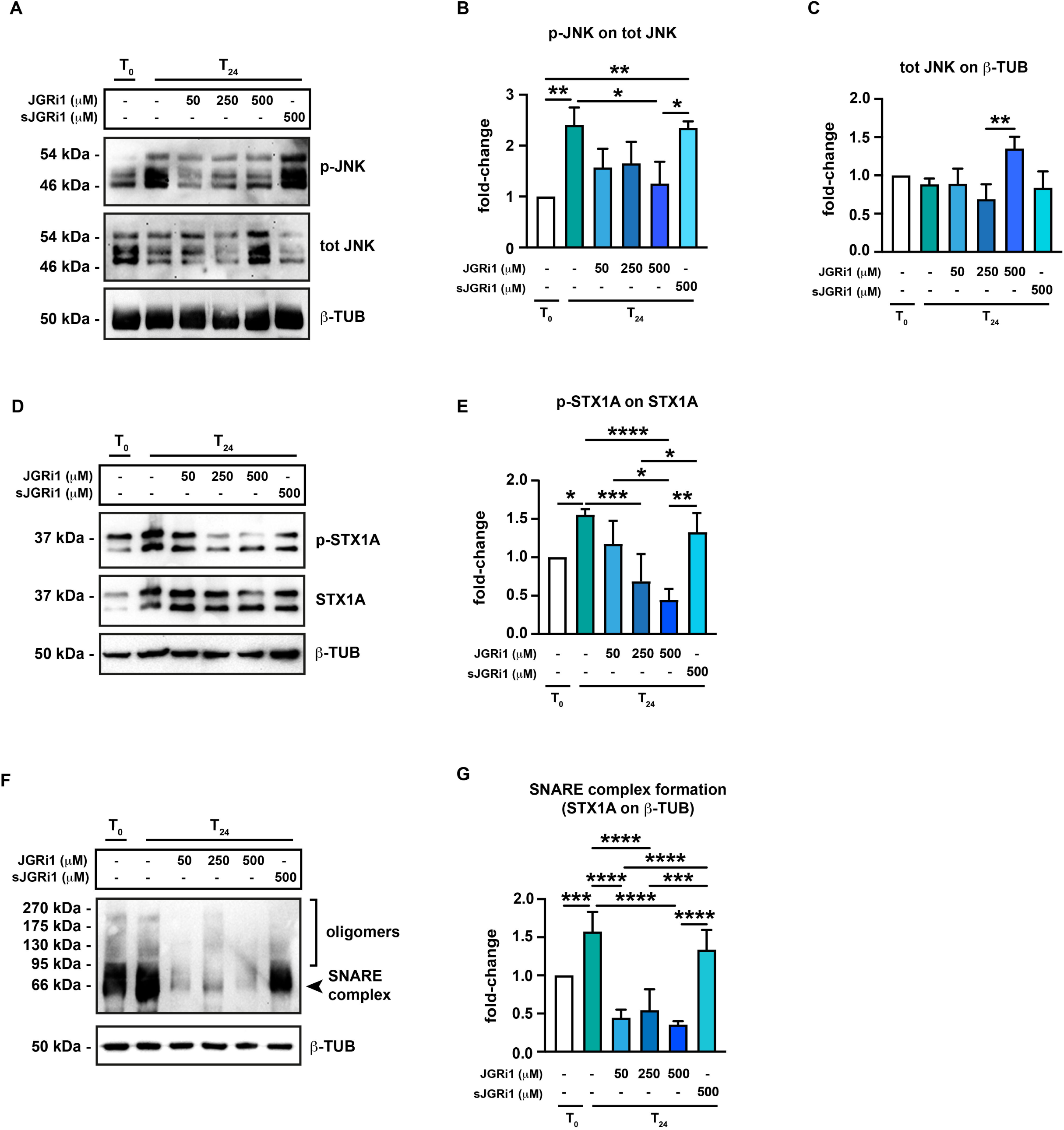
Effect of JGRi1 on the phosphorylation of JNK2 and STX1A and on SNARE complex formation in evONC model. **(A)** Representative western blot showing the phosphorylation of JNK (p-JNK). Samples were blotted, then incubated with primary antibodies against p-JNK, total JNK and, ultimately, β-TUB (loading control). n = 5 independent experiments. **(B)** Densitometric analysis of p-JNK with respect to total JNK. n = 5 independent experiments. **(C)** Densitometric analysis of total JNK with respect to β-TUB. n = 5 independent experiments. **(D)** Representative western blot showing the phosphorylation of STX1A (p-STX1A). Samples were blotted, then incubated with a specific primary antibody against p-STX1A, then STX1A and, ultimately, β-TUB (loading control). n = 5 independent experiments. (**E)** Densitometric analysis of p-STX1A with respect to STX1A. n = 5 independent experiments. **(F)** Representative western blot showing SNARE complex formation in retinas from evONC model. Non-boiled protein samples were blotted, then incubated with primary antibodies against SNARE complex component STX1A and β-TUB (loading control). n = 6 independent experiments. **(G)** Densitometric analysis of STX1A with respect to β-TUB. n = 6 independent experiments. For reader’s convenience statistical significance only in comparison to T_24_ and between active JGRi1 and sJGRi1 is shown. **Bar plot caption:** Bar representing mean +/− S.D. Statistical analysis: One-way ANOVA, *post hoc* Tukey test, p < 0.05.

The release of glutamate entails the formation of Soluble N-ethylmaleimide-Sensitive Factor Attachment Proteins Receptor (SNARE) complexes which are multiprotein complexes that drive the fusion of neurotransmitter-storing vesicles to plasma membrane (Ramakrishnan et al., 2012; Rizo, 2018). Therefore, the rate of SNARE complex formation in the retina was assessed in the evONC model. Incubation with STX1A antibody revealed the presence of a band at around 66 kDa, corresponding to the core SNARE complex, plus a high molecular weight protein smear, which corresponds to its different oligomers (Fig. 8F), in line with previous reports (Tokumaru et al. 2001). The intensity of the signals increased at T_24_ indicating that evONC boosted SNARE complex formation, while the treatment with all the doses of JGRi1 significantly reduced signal intensity. Treatment with sJGRi1 did not produce any effect on SNARE complex formation (Fig. 8G-H). These data show that JGRi1 reduces the evONC-boosted formation of SNARE complexes.

### JGRi1 treatment reduced NMDA-induced RGC death in retinal wholemount preparations

The protective effect of JGRi1 was assessed also on NMDA-treated retina wholemounts. To do so, wholemounts were treated with 1 μM JGRi1 for 1 hour prior to NMDA stimulation. (Fig. 9A). Firstly, RGC viability was assessed by immunoblot for the BRN3A. As shown previously, at T_24_ BRN3A reduction was enhanced by treatment with 100 μM NMDA. However, treatment with 1 μM JGRi1 prior to NMDA was able to rescue the NMDA-enhanced reduction of BRN3A expression, bringing it back almost to control levels (Fig. 9B-C). Moreover, our previous data showed that at T_24_ 100 μM NMDA treatment produced an increase of GS expression. Treatment with 1 μM JGRi1 produced a significant reduction of GS levels in comparison only to NMDA-treated retinas, but not retinas at T_24_ (Fig. 9D-E).

**Figure 9.**
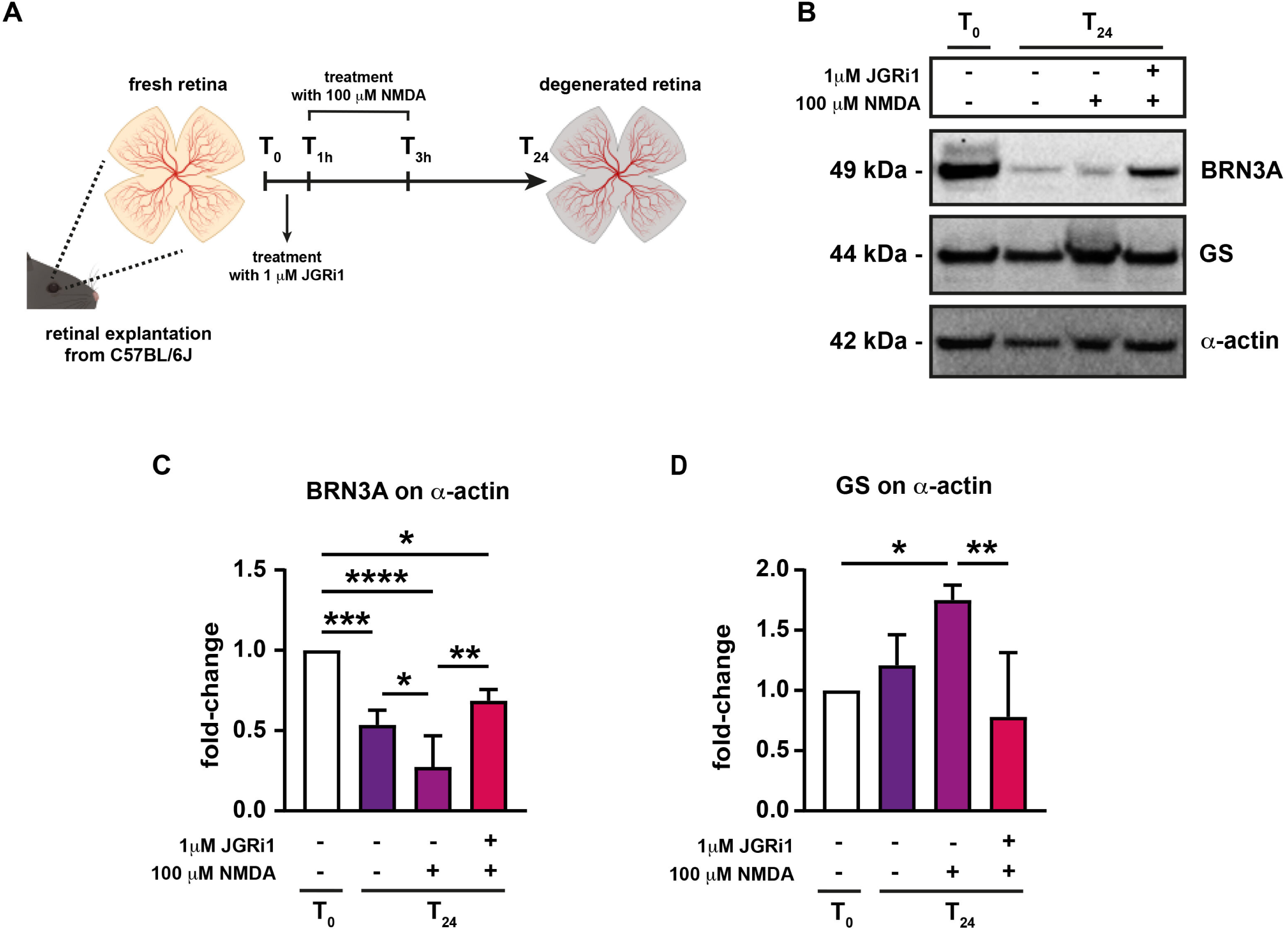
JGRi1 prevents NMDA-induced loss of RGCs and boost in GS expression in retina wholemounts. **(A)** Schematic representation of JGRi1 treatment on retina whole-mount preparations. **(B)** Representative western blot showing the expression of BRN3A and GS in wholemount retinas. Samples were blotted, then incubated with primary antibodies against BRN3A, GS and, ultimately, α-actin (loading control). n = 5 independent experiments. **(C)** Densitometric analysis of BRN3A with respect to α-actin. n = 5 independent experiments. **(D)** Densitometric analysis of glutamine synthetase with respect to α-actin. n = 5 independent experiments. **Bar plot caption:** Bar representing mean +/− S.D. Statistical analysis: One-way ANOVA, *post hoc* Tukey test, p < 0.05.

### Topical JGRi1 protects the GCL and reduces glutamate release in mouse NMDA-induced retinal degeneration model

As next goal, the protective effect of JGRi1 in the NMDA-induced retinal degeneration model was addressed. To do so, topical treatment with 250 μM of JGRi1, one drop/die, was started one day prior to NMDA injection, then kept on throughout the entirety of the follow-up post-injection. At the end, animals were sacrificed and eyeballs collected for the analyses. sJGRi1 was include as control (Fig. 10A). Firstly, the protection of JGRi1 on RGC was measured by performing immunofluorescence for the RGC marker NeuN and the apoptotic marker c-CASP3 (Fig. 10B), followed by quantification of NeuN-positive and c-CASP3-positive cells. Data showed that JGRi1 was able to rescue RGC viability upon NMDA injection (Fig. 10B-C) Conversely, c-CASP3 immunoreactivity, which increased during NMDA injection, was reduced by JGRi1 treatment (Fig. 10B-D). As expected, treatment with sJGRi1 had no effect on either RGC viability and apoptosis.

**Figure 10.**
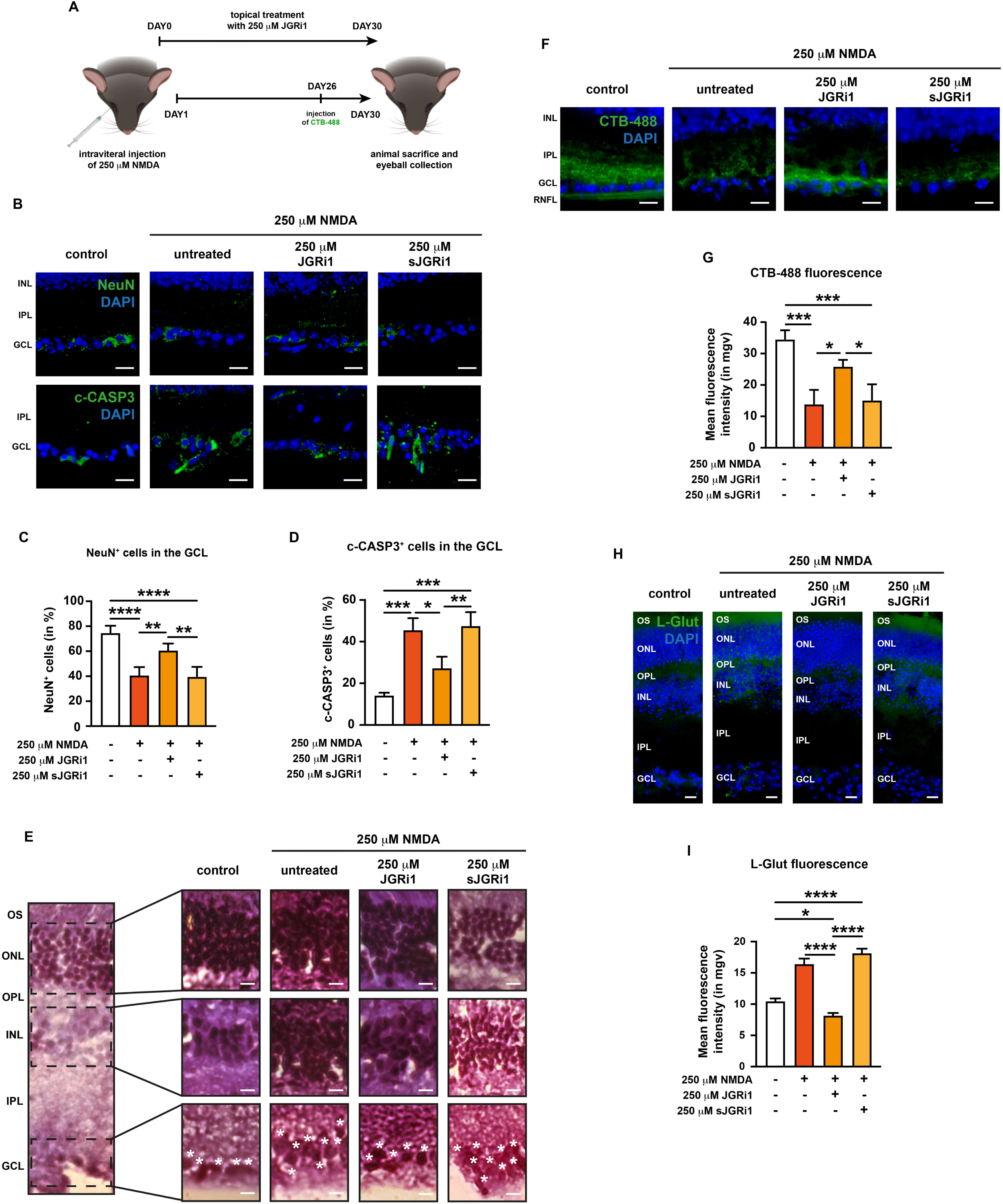
JGRi1 protects retinas from NMDA-induced degeneration. **(A)** Schematic representation of JGRi1 treatment. **(B)** Representative immunofluorescence for NeuN (upper row) and c-CASP3 (bottom row). Eyes were processed for immunofluorescence with specific antibody (green). n = 4 independent experiments. **(C)** Quantification of NeuN-positive cells. n = 4 independent experiments. **(D)** Quantification of c-CASP3-positive cells. n = 4 independent experiments. **(E)** Representative hematoxylin-eosin staining on retinal slices from NMDA-injected mice. Eyes were stained with hematoxylin, then counterstained with eosin. A sample retina is shown in the left part of the panel. For each nuclear layer an enlarged view is provided. White asterisks indicate RGCs. n = 3 independent experiments. **(F)** Representative immunofluorescence for CTB-488. n = 5 independent experiments. **(G)** Quantification of the mean fluorescence intensity of CTB-488. n = 4 independent experiments. **(H)** Representative immunofluorescence for L-glut. Eyes were processed for immunofluorescence with a L-glut-specific antibody (green). n = 3 independent experiments. **(B)** Quantification of the mean fluorescence intensity of L-glut. n = 3 independent experiments. **Immunofluorescence caption:** OS, outer segment; ONL, outer nuclear layer; OPL, outer plexiform layer; INL, inner nuclear layer; IPL, inner plexiform layer; GCL, ganglion cell layer. 40X magnification. Scale bar 10 μM. **Bar plot caption:** Bar representing mean +/− S.D. Statistical analysis: One-way ANOVA, *post hoc* Tukey test, p < 0.05.

Hematoxylin/Eosin staining was carried out to evaluate the retinal cytoarchitecture in the experimental conditions (Fig. 10E). The results show that NMDA injection produced a disruption of the three nuclear layers of the retina, with a major effect on the INL and GCL in which cell nuclei lost their physiological tissue compactness, as previously reported (Lambuk et al., 2019; Ishimaru et al., 2017). On the other end, 250 μM JGRi1 seemed to restore the tissue architecture of the GCL, while sJGRi1 failed to do so (Fig. 10E – white asterisks).

The protective effect of JGRi1 on NMDA-induced RGC damage was assessed by uptake and transport of the neural tracer CTB-488. The analysis showed that NMDA alone and with sJGRi1 treatment diminished CTB uptake and transport in RGCs. In contrast, treatment with JGRi1 was able to prevent at least in part this NMDA-induced decrease, suggesting that the peptide preserves the functional integrity and health of RGCs in the presence of excitotoxic stress (Fig. 10F-G).

Lastly, to evaluate L-glutamate levels in retinal tissue from the NMDA-injected eyes the L-glutamate immunofluorescence was measured. While NMDA injection led to a higher glutamate immunoreactivity in the murine retina, especially between the INL and ONL, concomitant treatment with 250 μM JGRi1 caused its reduction, even below control levels (Fig. 10H-I). sJGRi1, on the other hand, produced no effect on L-glutamate immunoreactivity. These data show that JGRi1 can protect RGC from NMDA-induced degeneration and prevent glutamate spillover.

### JGRi1 reduces the NMDA-induced expression of JNK2 and STX1A and prevents their interaction

The behavior of STX1A and JNK2 in response to the different treatments was then examined by co-immunofluorescence. As already shown in Fig. 3K-L, NMDA injection resulted in more JNK2 and STX1A expression in the retina of injected C57BL/6J mice (Fig. 11A). Treatment with JGRi1, but not sJGRi1, was able to reduce JNK2 immunoreactivity in the inner retina, which remained detectable in the GCL and slightly in the ONL as well. As expected, sJGRi1 did not prevent the NMDA-induced increase in JNK2 expression in the inner retina (Fig. 11A-B). Interestingly, differently from what we have seen in the evONC model, here JGRi1 was able to prevent the NMDA-induced increase of STX1A immunoreactivity, which was subsequently confirmed by the quantification of the fluorescence intensity in the retina (Fig. 11A-C). To address the effect of the JGRi1 on the JNK2-STX1A interaction we employed two different approaches. The first approach consisted in running correlation analysis between the two fluorescent signals to obtain the Pearson’s score. JNK2 and STX1A showed a higher yet not significant Pearson’s score upon NMDA injection, while treatment with JGRi1, but not sJGRi1, led to a significant decrease (Fig. 11D). The second approach consisted in performing proximity ligation assay (PLA), which allows to visualize the interaction between the two proteins as red puncta. In all tested conditions, PLA puncta were detectable mainly in the GCL and in the IPL (Fig. 11E). The number of PLA puncta increased in the slices of NMDA-injected mice, while it was reduced by treatment with JGRi1, but not sJGRi1 (Fig. 11F).

**Figure 11.**
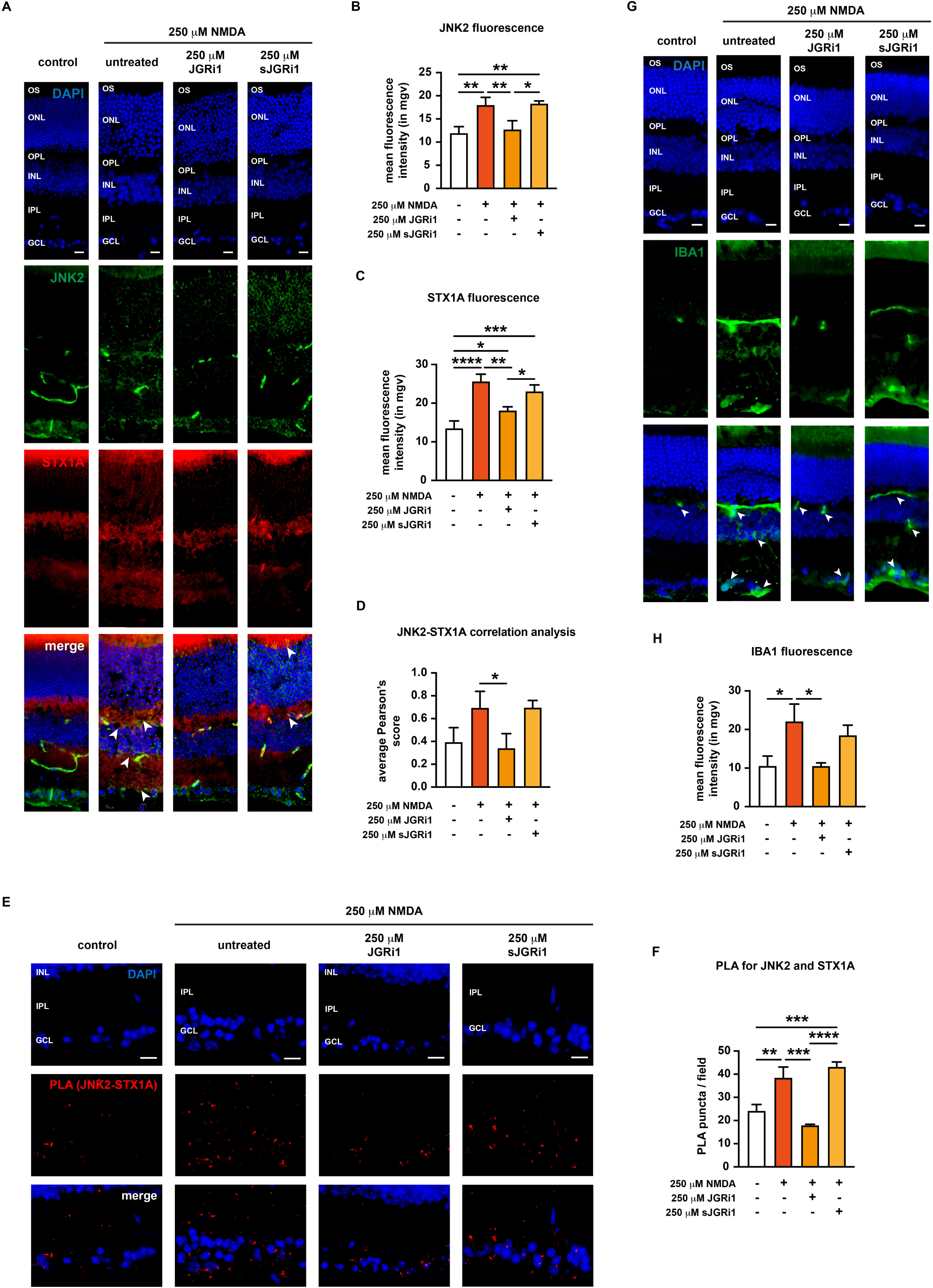
JGRi1 reduces the NMDA-induced expression of JNK2 and STX1A, reduces their interaction and microglial infiltration. **(A)** Representative co-immunofluorescence for JNK2 and STX1A. Eyes were processed for immunofluorescence with a JNK2 specific antibody (green) and a STX1A specific antibody (red). White arrowheads in the merge column indicate areas where JNK2 and STX1A signals strongly co-localize. n = 4 independent experiments. **(B)** Quantification of the mean fluorescence intensity of JNK2. n = 4 independent experiments. **(C)** Quantification of the mean fluorescence intensity of STX1A. n = 4 independent experiments. **(D)** Correlation analysis between JNK2 and STX1A signals. Pearson’s scores were calculated per each image from (A). n = 4 independent experiments. **(E)** Proximity ligation assay (PLA) for JNK2 and STX1A. Eyes were processed for PLA according to manufacturer’s instructions. Red puncta indicate JNK2-STX1A interactions. n = 4 independent experiments. **(F)** Quantification of PLA puncta per field. n = 3 independent experiments. **(G)** Representative immunofluorescence for IBA1. Eyes were processed for immunofluorescence with a IBA1-specific antibody (green). White arrowheads in the merge column indicate areas strongly immunoreactive to IBA1. n = 5 independent experiments. **(B)** Quantification of the mean fluorescence intensity of IBA1. n = 5 independent experiments. **Immunofluorescence caption:** OS, outer segment; ONL, outer nuclear layer; OPL, outer plexiform layer; INL, inner nuclear layer; IPL, inner plexiform layer; GCL, ganglion cell layer. 40X magnification. Scale bar 10 μM. **Bar plot caption:** Bar representing mean +/− S.D. Statistical analysis: One-way ANOVA, *post hoc* Tukey test, p < 0.05.

### JGRi1 reduces the NMDA-induced microglial infiltration in the retina

Microglial activation is an important contributor to RGC degeneration in different models of NMDA-induced retinal injury (Takeda et al., 2018; Sato et al., 2021). On this premise, the potential effect of the peptide on microglial reactivity in the retina of NMDA-injected mice was evaluated. Upon NMDA injection, a boost in IBA1 immunoreactivity was noticed throughout the entire retinal tissue, except the ONL (Fig. 11G). Surprisingly, treatment with JGRi1 was able to reduce IBA1 immunoreactivity, although some signal was still detectable in the OS, the IPL and the GCL, while no reduction was seen upon treatment with sJGRi1 (Fig. 11G-H).

## DISCUSSION

The presence of glutamate imbalance in RDs is well-established in animal models (McKinnon et al., 2009; Tsai et al., 2024; Mohamad et al., 2021; Kim et al., 2000), suggesting that glutamate accumulation is heavily involved in their pathogenesis. However, the strict relationship between glutamate spillover and RDs is not completely demonstrated in humans. Potentially, this is due to the fact that the increase of glutamate as well as the enzymes involved in glutamate metabolism are spatially restricted to the synapses, and in the retina, mainly in those between photoreceptors and bipolar cells and bipolar cells and RGCs. This fact makes extremely tricky the detection of glutamate elevation (Boccuni e Fairless 2022), but this phenomenon is still extremely harmful for the neuronal network functioning. Indeed, the toxicity of glutamate at synaptic level is well-documented in several neurological diseases (Hynd et al., 2004; Iovino et al., 2020; Van Den Bosch et al., 2006). In line with this, increasing evidence is showing off glutamate imbalance in RDs also for humans. A recent human retina atlas study pointed out at glutamate receptor and glutamate metabolism genes as risk factors for the onset of RDs (Yan et al., 2020). Proteins of the Excitatory Amino Acid Transporter (EAAT) family, like EAAT1 and EAAT2, which remove glutamate from the synaptic cleft, have been strongly linked to glaucoma since a recent study found two loss-of-function variants in the EAAT1 in patients suffering from glaucoma (Yanagisawa et al., 2020). Ultimately, studies carried out on retinas from retinitis pigmentosa patients highlighted extensive negative plasticity events, leading to the loss of the metabolic signature of Müller cell with subsequent glutamate accumulation (Jones et al., 2016).

The accumulation of glutamate inside the retina, in turn, causes an aberrant activation of downstream signaling cascades, including the ones involving NMDARs. NMDARs are extremely permeable to calcium ions; therefore, their overactivation will cause a massive calcium influx into the postsynaptic cell, culminating in the activation of a series of calcium-dependent apoptotic enzymes and, ultimately, in cell death. Such events go by the name of glutamate-mediated excitotoxicity. In accordance with the idea that excitotoxicity plays a role in the pathogenesis of RDs, anti-NMDA drugs, such as Riluzole, Resveratrol and Memantine, showed encouraging results in animal models for RDs, especially glaucoma (Esteruelas et al., 2022; Pirhan et al., 2016; Stefaniak-Clark et al., 2024; Atorf et al., 2013). However, despite their success in the pre-clinical phase, currently, there are no anti-NMDA drugs used in the clinical practice for the treatment of RDs. Memantine, which is one of the main anti-NMDA drugs available on the market, has failed in clinical trials for the treatment of glaucoma (Weinreb et al., 2018; Osborne, 2009). The failure of memantine is potentially due to several reasons: firstly, memantine is an uncompetitive NMDA receptor antagonist which turns off the NMDAR function, leading to severe side effects arising from the suppression of the glutamatergic tone in both the retina and the CNS; secondly, the oral administration does not guarantee a sufficient accumulation of the drug within the retina and the dosages cannot be increased because of CNS-related side effects; lastly, other possible routes of administration, such as intraocular injections, may cause severe discomfort to the patients. Despite all these downsides, targeting the NMDAR cascade still represents a valid approach for the treatment of RDs. Indeed, oral dextromethorphan, which acts primarily as an inhibitor of microglial activation, but also possesses anti-NMDA activity, was well-tolerated by patients and reached all primary endpoints in phase I/II clinical trials for diabetic macular edema (Valent et al., 2018).

Our previous work identified a novel NMDA-related mechanism called nPING, according to which the overactivation of presynaptic NMDARs causes the JNK2-dependent phosphorylation of STX1A (Nisticò et al., 2015), with a following enhancement of vesicle release and, finally, glutamate spillover (Nisticò et al., 2015; Marcelli et al., 2019). Our data showed that both in the evONC model and the intravitreal NMDA injection model there was an upregulation of JNK2 and STX1A. We also show that NMDA alone is less effective than axotomy (with or without NMDA) at inducing JNK2-STX1A interaction. This may indicate that the nPING mechanism may be activated by mechanical injury or stress to RGC axons. This would then “set the stage” for subsequent excitotoxic injury, thus pointing out to a potential biphasic mechanism for RGC degeneration, especially in glaucoma. Indeed, we speculate that intraocular pressure elevation in glaucoma may trigger nPING, thus fostering a subsequent excitotoxic damage.

The increased JNK2-STX1A interaction was accompanied also by and increase in retinal glutamate immunoreactivity. The higher glutamate levels found in both models are in line with previous reports, as well as the higher JNK2 levels (Hassanzadeh et al., 2022). Indeed, works from other groups showed that JNK2 is specifically activated and contributes to RCG death in both a retinal axonal injury-induced model and in an intravitreal endotelin-1 injection model (Kodati et al., 2021; Fernandes et al., 2012), although another paper suggested that JNK2 may protect RGC from glaucomatous degeneration in ocular hypertensive DBA/2J mice (Harder et al., 2018). Interestingly, to date, there were no reports about changes in the expression of STX1A levels in specific RDs. In addition, our data from both Pearson’s analysis and PLA suggested that upon retinal damage there is a higher interaction between JNK2 and STX1A. This evidence, combined with the increased glutamate immunoreactivity found in both the evONC and the NMDA injection model, suggests that further research is needed to understand the full extent of role of nPING and its potential implications for the pathogenesis of RDs. Moreover, since excitotoxicity is a mechanism shared by many neurological disorders, nPING possesses a high translatability potential, making it a valid molecular pathway to be targeted for the treatment of CNS diseases.

To selectively disrupt nPING, we designed molecular docking analyses a small peptide sequence which has then been linked to the highly-permeable sequence of the Tat protein of HIV-1, resulting in a cell-permeable peptide called “JGRi1” (Marcelli et al., 2019). The usage of a cell-permeable peptide represents opens up new therapeutic avenues by allowing the topical administration of the compound directly onto the eye, thus contrasting the two major downsides of the oral administration, namely the poor retinal accumulation and the subsequent higher risk of side effects due to drug overdosage. Indeed, our data produced using fluorescently-tagged JGRi1showed that the peptide was able to reach the retina upon both *ex vivo* and *in vivo* treatments.

RGC are particularly vulnerable to excitotoxicity, although with some differences between the various subtypes. Indeed, αRGCs are the most resistant type of RGCs to NMDA excitotoxicity, while the J-RGCs are the most sensitive cells (Christensen et al., 2019). Our data clearly show that JGRi1 can actually prevent RGC death to some extent. Such evidence represents a landmark step, as RGC degeneration underlies several conditions which give rise to significant visual compromise, including glaucoma, hereditary optic neuropathies, ischemic optic neuropathies, and demyelinating disease (Khatib e Martin, 2017). Further studies will be needed in order to determine which RGC subtype benefits the most from JGRi1 treatment.

Moreover, JGRi1 successfully disrupted the JNK2-STX1A interaction in both evONC and the NMDA injection models, as shown by the Pearson’s and PLA analyses, and reduced glutamate levels in treated retinas. To our knowledge, this study provides the first demonstration that a presynaptic protein–protein interaction, namely the JNK2–STX1A complex, can be effectively disrupted in vivo by a cell-permeable peptide, highlighting a novel therapeutic paradigm for targeting excitotoxic neurotransmission. Indeed, JGRi1 was able to preserve retinal cytoarchitecture as well as RGC viability. RGCs degeneration underpins many RDs (Khatib e Martin, 2017). JGRi1 was able to preserve their viability in all the tested models, as shown by immunofluorescences for RBPMS and NeuN and by western blot for BRN3A.

In the evONC model, JGRi1 prevented the boost in JNK and STX1A phosphorylation, in line with our previous reports from SH-SY5Y cells and murine brain-derived synaptosomal fractions (Marcelli et al., 2019; Cimino e Feligioni, 2024). Worth of notice, JNK2 expression levels remained significantly higher than the control upon treatment with all tested dosages of JGRi1. Such result may be in line with the idea that JNK2 may protect RGC from degeneration to some extent, in line with previous reports (Harder et al., 2018).

As mentioned previously, SNARE complexes are those responsible for neurotransmitter release and their assembly is strongly influenced by STX1A phosphorylation ((Dubois et al., 2002; Foster et al., 1998). Our measurement of SNARE complex formation in the evONC retina showed that JGRi1 led to an almost complete abrogation of SNARE complex assembly in the retina. Such evidence, combined with the fact that higher dosages of JGRi1 dramatically reduced retinal glutamate levels, suggests that high dosages of JGRi1 may be detrimental for normal retinal physiology, as they shut down retinal neurotransmission. Therefore, further studies are needed in order to figure out the effect of SNARE complex reduction in the murine retina and to define the best dosage regimen of JGRi1 that does not shut down retinal neurotransmission completely.

GS is one of the most important enzymes involved in retinal glutamate metabolism, as it is highly expressed in the Müller glia, where it coverts glutamate to glutamine (Bringmann et al. 2013). Interestingly, our data from retinal wholemount preparations showed an upregulation of GS at 24 hours post-mounting, indirectly indicating that glutamate level in the synaptic cleft was high. GS levels tented to be even more enhanced by the treatment with NMDA, while they were brought back to control levels by the treatment with JGRi1. Such finding may seem quite in contrast with what has already been reported, as GS levels were found to be reduced in the retina of patients suffering from geographic atrophy (Naik et al., 2025). However, a previous paper showed that after retinal ischemia GS can be upregulated for a short period of time before its levels start to decay. Therefore, since our retinal wholemounts were cultured for 24 hours, we believe that the higher GS levels may represent an early event occurring in order to prevent NMDA-induced damage (Wang et al., 2024).

Lastly, microglial infiltration has been reported in different models of NMDA-induced retinal injury (Takeda et al., 2018; Sato et al., 2021). Our data clearly showed that JGRi1 was able to reduce microglial infiltration inside the retinas from NMDA-injected mice. A possible explanation to this may lay in the fact that JNK2 promotes microglial activation and the release of pro-inflammatory mediators (Waetzig et al., 2005). Therefore, we speculate that JGRi1 by inhibiting JNK2 may also reduce the degree of microglial infiltration within the tissue. However, further studies will be needed in order to address this specific topic.

Taken together, our findings indicate that the nPING mechanism plays a critical role in retinal degeneration. Future investigations on JGRi1 are warranted to corroborate our hypothesis about mechanical injuries as potential triggers to nPING, to optimize JGRi1 formulation and evaluate its potential across multiple therapeutic indications, encompassing both RDs and CNS diseases. We believe that this work lays the groundwork for the development of a new generation of pharmacological strategies specifically targeting excitotoxic presynaptic mechanisms in neurodegeneration.

## MATERIALS & METHODS

### Animals

C57BL/6J mice (male, 25–30 g, 4-5 weeks) were rendered and housed in a temperature- and humidity-controlled condition on a 12:12 light-dark cycle with ad libitum food. At the end of treatments, animals were euthanized via beheading or transcranial perfusion with heparin-containing saline followed by 4% paraformaldehyde. Furthermore, since the glutamate synthesis and release are circadian-dependent, animals from all experimental groups were sacrificed at the same daytime.

### *Ex vivo* permeability assay

To assess JGRi1 permeability *ex vivo*, eyeballs from euthanized animals were collected and immersed into a balanced salt solution (BSS) buffer (137 mM NaCl, 5.4 mM KCl, 1.8 mM CaCl_2_•2H_2_O, 0.98 mM MgCl_2_•6H_2_O, 8.1 mM Na_2_HPO_4_•2H_2_O, 1 mM K_2_HPO_4_, 5.5 mM D-glucose, 4.2 mM NaHCO_3_, pH 7.4), then incubated for 1 hour at 37°C with different concentrations (50 μM, 250 μM, 500 μM) of a fluorescently labeled version of our JGRi1 peptide (F-JGRi1). A version of F-JGRi1 devoid of the Tat sequence, which grants its cellular permeability (ΔTat-F-JGRi1), was included as additional control. Afterwards, eyeballs were washed three times with BSS to remove excess of the peptide, then processed according to experimental needs.

### *In vivo* permeability assay

For the *in vivo* permeability assay of JGRi1, animals were treated with a single drop per day of F-JGRi1 at different concentrations (50 μM, 250 μM, 500 μM) dissolved into BSS buffer. A total volume of 5 μL of JGRi1 solution applied directly onto the cornea. Treated animals were kept immobilized and under observation for 1 minute to ensure proper peptide penetration within the eye. At the end of treatments, animals were sacrificed, eyeballs were collected and processed accordingly. ΔTat-F-JGRi1 was also included as additional control.

### *Ex vivo* optic nerve cut (evONC)

Mice were either left untreated or treated for 6 days with a single drop per day of JGRi1 dissolved into a balanced saline solution (BSS) buffer at different concentrations (50 μM, 250 μM, 500 μM). A total volume of 5 μL of JGRi1 solution applied directly onto the cornea. Treated animals were kept immobilized and under observation for 1 minute to ensure proper peptide penetration within the eye. Animals were then euthanized via beheading. A scrambled version of the peptide (sJGRi1) at a concentration of 500 μM was included as control. Subsequently, the optic nerve was cut and eyes were harvested according to our previously established protocol (Hassanzadeh et al., 2022; Buccarello et al., 2021). The extracted eyes were kept in phosphate buffered salt (PBS; Capricorn Scientifics, Germany) at 4°C for 24 hours, then the retinas were dissected and processed accordingly.

### Retinal wholemounts

Eye bulbs from C57BL/6J mice were explanted and put into a Petri dish in presence of DMEM without fetal bovine serum (FBS) supplementation. Using a stereomicroscope placed under a cell-culture hood, retinas were isolated using surgical forceps, then cut into quarters using surgical scissors starting from the retinal periphery all the way down to the optic nerve emergence. Afterwards, explants were cultured in 6-well plates containing either fixed or removable inserts (Euroclone, ET3006). 1 mL of DMEM, supplemented with 1:100 penicillin/streptomycin (Lonza, Switzerland) and 10% v/v FBS (Capricorn Scientific, Germany) was added put inside the well, underneath the insert. Explants were placed on the insert with the ganglion cell layer facing upward, in order to improve photoreceptor survival through contact with the medium beneath the insert. To maintain hydration, an additional 100 μL of medium was applied directly onto the insert, ensuring that the explant remained moist without floating. Afterwards, retinas were incubated for 1 hour at 37°C, treated with 100 μM NMDA (Merck, Germany) for 2 hours, then reverted to control medium. At 24 hours post-mounting, retinas were processed accordingly. JGRi1 or sJGRi1 at the concentration of 1 μM were applied 1 hour prior to NMDA treatment.

### NMDA Injection Model

Retinal excitotoxicity was induced unilaterally in adult male C57BL/6J mice (n=18) by intravitreal injection (2 μL) of 250 μM NMDA. Mice were anesthetized using isoflurane gas at 4–5% for induction in an enclosed chamber, then maintained at 2–3% via nose cone during the procedure. Prior to injection, a few drops of proparacaine (0.5% solution) were applied to both eyes to provide local numbing. Unilateral intravitreal injections of NMDA were to the left eye and equivalent volume of saline (control) was injected into the vitreous cavity of the mouse using a 33-gauge needle attached to a Hamilton syringe. To minimize risk of postoperative infection, Tobramycin Ophthalmic Solution (0.3%) was applied topically to the cornea. For pharmacological treatment, mice were randomly assigned to one of three groups: active peptide treatment (n = 6), scrambled peptide control (n = 6), or an untreated control (n = 6). Mice were treated with either an active form of JGRi1 or sJGRi1. Mice were treated daily with 4 µL of 250 µM JGRi1 or 250 µM sJGRi1 applied directly to the cornea as a topical eye drop. Prior to drug treatments, mice were acclimated to scuffing and ocular drop administration. Drug administration began 24 hours prior to NMDA injection and continued throughout the 4-week study period.

Since anterograde axonal transport is a critical indicator of RGC integrity and visual pathway connectivity, we employed Cholera Toxin Subunit B (CTB), well-established neural tracer that is actively transported along RGC axons. Five days prior to sacrifice, all mice received bilateral intravitreal injections of 1 µL of CTB conjugated to Alexa Fluor 488 (CTB-488; Crish et al., 2010; Echevarria et al., 2017; Wareham et al., 2021; Fischer et al., 2019).

### Preparation of Retinal Lysates

After enucleating the eyes, the retinas immediately were extracted and lysated in 100 μl of RIPA buffer (50 mM Tris-HCl pH 7.4, 150 mM NaCl, 1% v/v NP-40, 0.1% w/v sodium-dodecyl-sulphate (SDS), 0.5% w/v sodium deoxycolate, 1 mM ethylenediaminetetraacetic acid (EDTA)) supplemented with a complete set of protease inhibitors (Complete, Roche Diagnostics, Basel, Switzerland) and phosphatase inhibitors (Sigma, St. Louis, MO). Samples were then sonicated, and the homogenates were placed on ice for 30 min to allow protein solubilization. Then, they were centrifuged at 10000 rpm for 10 min, and subsequently, the supernatant was collected and stored at −80°C until needed. Samples concentration was measured by quantifying the A_280_ using a nanodrop system (ThermoFisher).

For retina wholemounts, proteins were extracted using a glass homogenizer containing an extraction solution (50 mM Tris HCl pH 7.5, Triton X 100%, 20% SDS, 0.5 M EDTA), supplemented with a protease and phosphatase inhibitor cocktail (Thermo Scientific, US). The retinas were left on ice for 20 minutes to allow the extraction solution to act and finally centrifuged at 13200 rpm at 4°C. Subsequently, the supernatant containing the proteins was collected.

### Western Blot

For each sample, 50 μg of protein extract supplemented with 1X Laemmli buffer with the addiction of 2.5% β-mercaptoethanol were boiled at 95°C for 5 minutes, then separated on 10-15% SDS polyacrylamide gel electrophoresis (SDS-PAGE) and transferred to PVDF membranes. Afterwards, the membranes were blocked in 5% non-fat milk powder dissolved into Tris-buffered saline (TBS) supplemented with 0.1% v/v Tween 20 (TBS-T) for 1 hour at room temperature. Afterwards, membranes were incubated overnight at 4°C with the following primary antibodies diluted either in the same blocking solution or 3% bovine serum albumin (BSA) dissolved in TBS-T: anti-RBPMS (ab15210, Abcam, UK); anti-c-CASP3 (9661, Cell Signaling, US); anti-JNK2 (sc-271133, Santa Cruz, US); anti-STX1A (S0664, Merck, Germany); anti-p-STX1A (Ser^14^) (CSB-PA053965, Cusabio, US); anti-STX1A (110302, Synaptic System, Germany); anti-p-JNK (9251, Cell Signaling, US); anti-JNK (9252, Cell Signaling, US); anti-BRN3A (sc-8429, Santa Cruz, US); anti-GS (ab64613, Abcam, UK). To develop the blots, horseradish peroxidase-conjugated secondary antibodies (anti-mouse or anti-rabbit, 1: 10000, Bio-Rad, US) were utilized, and the immunoreactive bands were visualized by exposure to the ECL chemiluminescence system (Merck, Germany). Either β-tubulin (ab18207, Abcam, UK) or α-actin (sc-1616, Santa Cruz, US) were used as the loading control for quantification. Western blots were quantified by densitometry using ImageJ software. Protein expression was expressed as fold-change and obtained by dividing the band density of the sample by that of the loading control. The activation of both JNK and STX1A was expressed as the ratio of the band intensity of phosphorylated form to the total protein.

### SNARE complex assay

The modulation SNARE complex assembly by JGRi1 was evaluated via native PAGE. Briefly, samples loaded using a non-denaturizing loading buffer (0.5 M Tris-HCl pH 6.8, 15% v/v glycerol and 1% v/v bromophenol blue). Samples were then left non-boiled to preserve protein-protein interaction, separated onto polyacrylamide gels, then blotted onto nitrocellulose membranes. The immunoreactive bands identified by using an antibody against STX1A, which is a core component of SNARE complexes. Bands starting from approximately 66 kDa were quantified as an indicator of the SNARE complex assembly upon treatment. β-tubulin was used as the loading control for quantification. the rate of SNARE complex formation was expressed as fold-change and obtained by dividing the band density of the sample by that of the loading control.

### Hematoxylin/Eosin staining

The enucleated eyes were fixed in 4% paraformaldehyde solution overnight at 4°C. Then, they were incubated in a 30% sucrose-in-PBS solution overnight, and finally, they were embedded in an optimal cutting temperature (OCT, Sigma, St. Louis, MO, USA) compound. Eyes were cut at a thickness of 20 μm. For reliability, the sections containing the optic nerve were utilized, and in each eye, at least five discontinuous sections were analyzed. For histological examination cryosections were stained in Mayer’s Hematoxylin Solution (Merck, Germany), rinsed in tap water, then counterstained using Eosin Y alcoholic solution (Merck, Germany). Sections were mounted on coverslips with Eukitt (Merck, Germany), and observed under a light-transmission microscope (Nikon) with 40X magnification.

### Immunofluorescence

Retinal cryosection were permeabilized using 0.5% Tryton-X (Merck, Germany) in PBS (Capricorn Scientifics, Germany) solution for 5 minutes. Then, blocked (0.1 M glycine, 2% w/v bovine serum albumin, 2% v/v fetal calf serum, 0.1% v/v Tryton-X) for 1 hour to avoid non-specific protein interactions. The primary antibody was diluted in the same blocking solution and held overnight at 4°C. The following primary antibodies were used: anti-RBPMS (ab15210, Abcam, UK); anti-c-CASP3 (9661, Cell Signaling, US); anti-L-Glutamate antibody (ab9440, Abcam, UK); anti-JNK2 (sc-827, Santa Cruz, US and GTX107477, GeneTex, US); anti-STX1A (S0664, Merck, Germany); anti-STX1A (110302, Synaptic System, Germany); anti-IBA1 (019-19741, Fujifilm Wako, JP). The secondary antibodies were Alexa Fluor® 488 and Alexa Fluor® 594 conjugated-anti-rabbit or anti-mouse IgG (Invitrogen, US), dissolved in the same blocking solution at a 1:500 dilution for 1 hour. Slides were subsequently mounted on coverslips using a DAPI-containing mounting medium (Fluoromount, Invitrogen, US), and fluorescent images were acquired using a confocal laser scanning microscope at a 40X magnification (Olympus; Confocal System FV500). Fluorescence intensity was quantified by using ImageJ software. Fluorescence intensity was expressed as mean gray values (mgv) minus background signal calculated in least 5 different fields per condition. To calculate the number of RBPMS and c-CASP3-positive cells in the GCL, the total number of positive cells was divided by total number of cells in the GCL, then converted in percentage. At least 100 cells per condition were used for the analysis.

### Quantification of F-JGRi1 signal

Retinal cryosections were mounted on coverslips using a DAPI-containing mounting medium (Fluoromount, Invitrogen, US), and fluorescent images were acquired using a confocal laser scanning microscope at a 40X magnification (Olympus; Confocal System FV500). To measure F-JGRi1 signal intensity in single retinal layers, each layer was selected and fluorescence in that given layer was quantified by ImageJ. Fluorescence intensity was expressed as mean gray values (mgv) minus background signal, calculated in least 10 different fields per condition. To calculate the number of FITC-positive cells in the GCL, the total number of FITC-positive cells was divided by total number of cells in the GCL, then converted in percentage. At least 100 cells per condition were used for the analysis.

### Quantification of CTB-488 signal intensity

Retinal cryosections were mounted on coverslips using a DAPI-containing mounting medium (Fluoromount, Invitrogen, US), and fluorescent images were acquired using a confocal laser scanning microscope at a 40X magnification (Olympus; Confocal System FV500). Fluorescence intensity was quantified by using ImageJ software. Fluorescence intensity was expressed as mean gray values (mgv) minus background signal calculated in least 5 different fields per condition.

### Proximity Ligation Assay (PLA)

PLA reaction was carried out according to the manufacturer’s instructions (Merck, Germany). Briefly, samples were incubated with primary antibodies overnight at 4°C and the day after with the PLA PLUS and MINUS probes for 1 hour at 37°C. All antibodies and probes were diluted in the buffers provided in the kit. Subsequently, probe ligation was carried out by incubating samples for 30 minutes at 37°C, followed by an amplification step done by incubating samples for 100 minutes at 37°C. Slides were then mounted on coverslips using the DAPI-containing mounting medium (Merck, Germany). Fluorescent images were visualized by confocal microscopy at a wavelength of 594 nm with 60X magnification. The number of PLA puncta was calculated using ImageJ software. The number of puncta per field was obtained by dividing the total number of counted PLA puncta per the number of field (n = 10) used for the analysis.

### Pearson’s co-localization analysis

Co-localization between fluorescent signals in acquired images was expressed using Pearson’s score, which describes the correlation of the intensity distribution between two distinct fluorescent signals. The analysis was carried out using the “JACOP” plug-in available for the ImageJ software. At least 10 different fields per condition were utilized for the quantification.

### Statistical Analysis

Statistical analysis was carried out using the Graph Pad Prism 9 program. Pairwise comparisons were analyzed running an unpaired t-test. For multiple comparisons a One-way ANOVA, followed by Tukey’s *post hoc* test was used. Samples were as graphed as mean ± SD with, at least, a statistical significance given at p < 0.05.

## Cartoons

The cartoons shown in this paper were made using the open-source version of Biorender.

## Ethical statement

The use of all the animals employed in this work complied with the ARVO statement for the use of animals in ophthalmic and vision research. The study was approved by the Italian Ministry of Health (Permit Number 800-2023 PR) and by the IACUC committee of Wake Forest University Health Sciences (A23-061).

